# Asymmetric listening conditions perturb verbal and nonverbal coordination dynamics during conversation

**DOI:** 10.64898/2026.10.19.760093

**Authors:** Hanlu He, Axel Ahrens, Mario Medoni, Rita Martí Torra, Ivana Konvalinka

## Abstract

Human conversation depends on continuous coordination across verbal and nonverbal behaviour. Yet this coordination may become perturbed when one person experiences greater listening difficulty. Previous nonverbal joint action studies show that unequal task demands between partners can lead to complementary leader-follower dynamics, but whether similar patterns emerge during natural conversation remains unclear. We investigated how unequal listening conditions affect verbal and nonverbal coordination in 40 same-gender stranger dyads during three 10-minute face-to-face conversations. Each dyad participated in noise-free and asymmetric noise conditions, with one person exposed to speech-shaped noise at a time. Asymmetric noise reduced conversational flow and enjoyment, and shifted the balance of verbal contributions: noise-exposed participants took longer turns and contributed more speaking time, while their noise-free partners produced more backchannels. Interpersonal head-movement coordination became temporally asymmetric under noise, with movements of the noise-exposed participant more often preceding those of their partner. In contrast, interpersonal hand-movement coordination did not differ between conditions. Within individuals, hand coordination was stronger when participants themselves experienced noise than when their partner experienced noise. These findings demonstrate that perturbing one partner’s auditory access affects coordination within and between individuals, revealing how unequal interaction demands produce different coordination patterns across verbal and nonverbal behaviour.

## 1 Introduction

Human face-to-face conversation is one of the most fundamental forms of human social interaction, allowing people to exchange information, establish shared understanding, and form and maintain social relationships [1, 2]. Yet successful conversation depends on much more than the content of what is said. Interacting partners must coordinate their behaviour in real time, anticipating each other’s upcoming utterances and actions while continuously and mutually adapting when they speak, move, and respond to one another [3–5]. Rather than unfolding as a simple exchange of discrete messages, conversation thus relies on the ongoing alignment of perceptual, linguistic, and motor processes across interacting partners [6–9]. Therefore, conversation can be understood as a tightly coupled communicative process in which coordination emerges across multiple behavioural and cognitive levels [2, 10, 11].

At the core of this coordination lies turn-taking, an exceptionally efficient coordination mechanism that allows speakers to alternate speaking and listening roles with minimal delay. Turn transitions are typically characterised by near-seamless short gaps and occasional overlaps, reflecting rapid predictive processing and fine-grained temporal adaptation [12–15]. This temporal precision is not just a structural feature of dialogue but carries important social meaning: faster and more smoothly coordinated turn transitions are associated with greater perceived conversational flow, stronger affiliation, and increased feelings of social connection [16, 17]. Even subtle disruptions in timing can therefore influence the subjective experience of interaction.

Importantly, conversational coordination extends well beyond the verbal channel. Speakers also align their nonverbal behaviours such as head movements, gestures, and posture, forming a rich multimodal system of coordination [6, 18–22]. These bodily signals are closely coupled with speech production and comprehension, supporting processes such as turn regulation, feedback signaling, and shared attention [23–25]. For example, head movements often signal engagement or understanding, while gestures contribute to the structuring and emphasis of information [9, 26]. Coordination also extends to physiological processes. Speakers align their breathing with turn-taking, timing inhalations to speech initiation [27], and dyads can exhibit synchrony in autonomic activity such as heart rate, particularly during emotionally engaging interactions [28–30]. Together, these findings highlight that conversation is inherently multimodal, requiring continuous coordination across distinct but tightly coupled channels.

A large body of research has characterised conversational coordination as highly adaptive and flexible, with interlocutors continuously adjusting to each other’s moment-to-moment behaviour rather than relying on explicit planning [10, 11, 31, 32]. This adaptive alignment is reflected across linguistic, temporal, and bodily dimensions, including speech timing, lexical and syntactic alignment, and movement coordination [6, 20, 33]. This process is often assumed to be broadly reciprocal, with both interlocutors continuously adapting to one another and jointly regulating the temporal and multimodal dynamics of the interaction [2, 11, 34]. However, this assumption is rarely examined directly, despite the fact that everyday interaction often takes place under conditions of asymmetric sensory access. Background noise, spatial separation, or individual differences in hearing ability can all introduce unequal access to incoming information and disrupt coordination [12, 35–37].

Prior work has shown that when listening conditions are globally degraded (i.e., affecting all interacting individuals), people tend to compensate by increasing vocal effort, exaggerating articulatory cues, or enhancing bodily signaling [35–38]. Background noise can also disrupt collaborative information exchange, changing how strongly individuals rely on one another’s contributions during joint decision-making [39]. These findings suggest that shared auditory challenges affect both communicative behaviour and the flow of information between interacting individuals. However, when constraints are unevenly distributed, symmetric adaptation may no longer be sufficient to sustain efficient interaction. Insights from joint action research suggest that under such asymmetric conditions, leader-follower roles may emerge such that the participant with higher task constraints becomes less adaptive and more task focused, assuming the role of a leader, while the participant with the lower task constraints remains more adaptive, hence becoming the follower [40, 41]. This shift from mutual adaptation toward complementary, leader-follower dynamics has been observed in nonverbal coordination tasks and has been interpreted as an implicitly negotiated coordination solution to imbalance in task demands or informational access [40, 42].

Conversation can also be understood as a form of joint action that requires continuous coordination toward a shared communicative goal. Hence, a similar shift in coordination dynamics is likely to occur when there is asymmetry in sensory access. However, it is still unclear how this shift manifests in natural face-to-face conversation across verbal and nonverbal behaviour, and how it relates to subjective experience.

In this study, we investigate how asymmetric auditory access influences inter- and intra-personal coordination dynamics during natural conversation. We experimentally induced asymmetry in interaction demands by introducing speech-shaped noise to one member of a dyad at a time, while the other participant retained normal listening conditions. We recorded multimodal behavioural data, including speech, head and hand movements, physiological ECG signals (which are not analyzed as part of this paper), as well as subjective reports of conversational experience.

We investigated three main hypotheses. First, we expected asymmetric noise to negatively impact perceived conversational flow and enjoyment [35, 36]. Building on prior work linking temporal coordination to affiliation [16, 17], we further expected that faster responses, or shorter floor-transfer offsets, would predict higher enjoyment across dyads. Second, we expected asymmetric listening conditions to shift the balance of verbal contributions between partners. The noise-exposed participant was therefore expected to take a more dominant role in floor-holding (i.e., longer turns) due to increased cognitive load and reduced ability to monitor turn-taking cues in real time [43], while we expected the noise-free partner to increase short feedback signals such as backchannels, potentially to support grounding and maintain mutual understanding [23]. Third, we expected these asymmetries to also be reflected in nonverbal coordination. Based on prior non-verbal joint action literature [40, 44], we anticipated a shift toward more directional coupling of interpersonal head movement coordination, consistent with a leader–follower pattern, with the noise-exposed participant leading the interaction more. For hand movements, we explored whether coordination would increase under noise, motivated by evidence that acoustic challenge can enhance bodily coordination and co-speech gestural behaviour [45–47], potentially reflecting compensatory signaling when auditory information is degraded.

In addition to coordination between individuals, interaction also has consequences for how individuals coordinate their own behaviour. Recent work suggests that interpersonal coordination can influence intrapersonal coordination dynamics, showing that when people become mutually coordinated, their own intrapersonal bodily rhythms become decoupled [48] and self-regulation becomes weakened [49]. Hence, changes in conversational demands may not only affect how strongly individuals align with each other, but also how tightly their own movements or physiological signals are coordinated over time. Asymmetric listening condition in this case may alter the balance between self- and other-oriented processing [50, 51]. Specifically, increased processing demands may lead the noise-exposed participant to become more internally focused, which could be reflected in stronger within-person coordination. Meanwhile, the noise-free partner may adapt more overtly to the interactional demands and partially decouple from their habitual movement patterns.

Together, this study uses asymmetric noise as a controlled perturbation to examine how conversational coordination is reorganised when interaction partners face unequal perceptual constraints. By integrating verbal, nonverbal, and subjective measures, we examine how adaptation is distributed between interaction partners and across behavioural levels. This provides a framework for testing whether unequal interactional demands shift conversation away from reciprocal mutual adaptation toward more complementary patterns of coordination.

## 2 Methods

### 2.1 Participants and study design

Forty same-gender dyads (N = 80 participants; 19 female dyads, 21 male dyads) were recruited for the study. Participants were native Danish speakers with normal hearing and no reported communication difficulties.

Ages ranged from 18 to 35 years, with a maximum age difference of four years within each dyad. The mean age was 22.61 years (SD = 1.80) for male participants and 23.11 years (SD = 3.27) for female participants. All participants were strangers prior to the experiment.

Participants attended the laboratory in groups and were allowed to interact freely during the preparation phase (approximately 20 minutes). Following preparation, participants completed a 10-minute resting-state electrocardiogram (ECG) recording while seated in isolation.

After this baseline recording, dyads were seated in a conversation booth and engaged in three 10-minute unstructured conversations in Danish. Participants were informed that auditory noise might be presented through their headphones during the conversations, but they were not told when or to whom the noise would be delivered. They were explicitly instructed not to disclose whether they were experiencing noise and to maintain the conversation as naturally as possible.

The experiment comprised a Noise-Free condition and two asymmetric noise conditions in which spectrally speech-shaped noise was presented to one participant at a time. Thus, each participant experienced one Noise-Self, one Noise-Other, and one Noise-Free interaction, with condition order counterbalanced across dyads (Figure 1).

**Figure 1.**
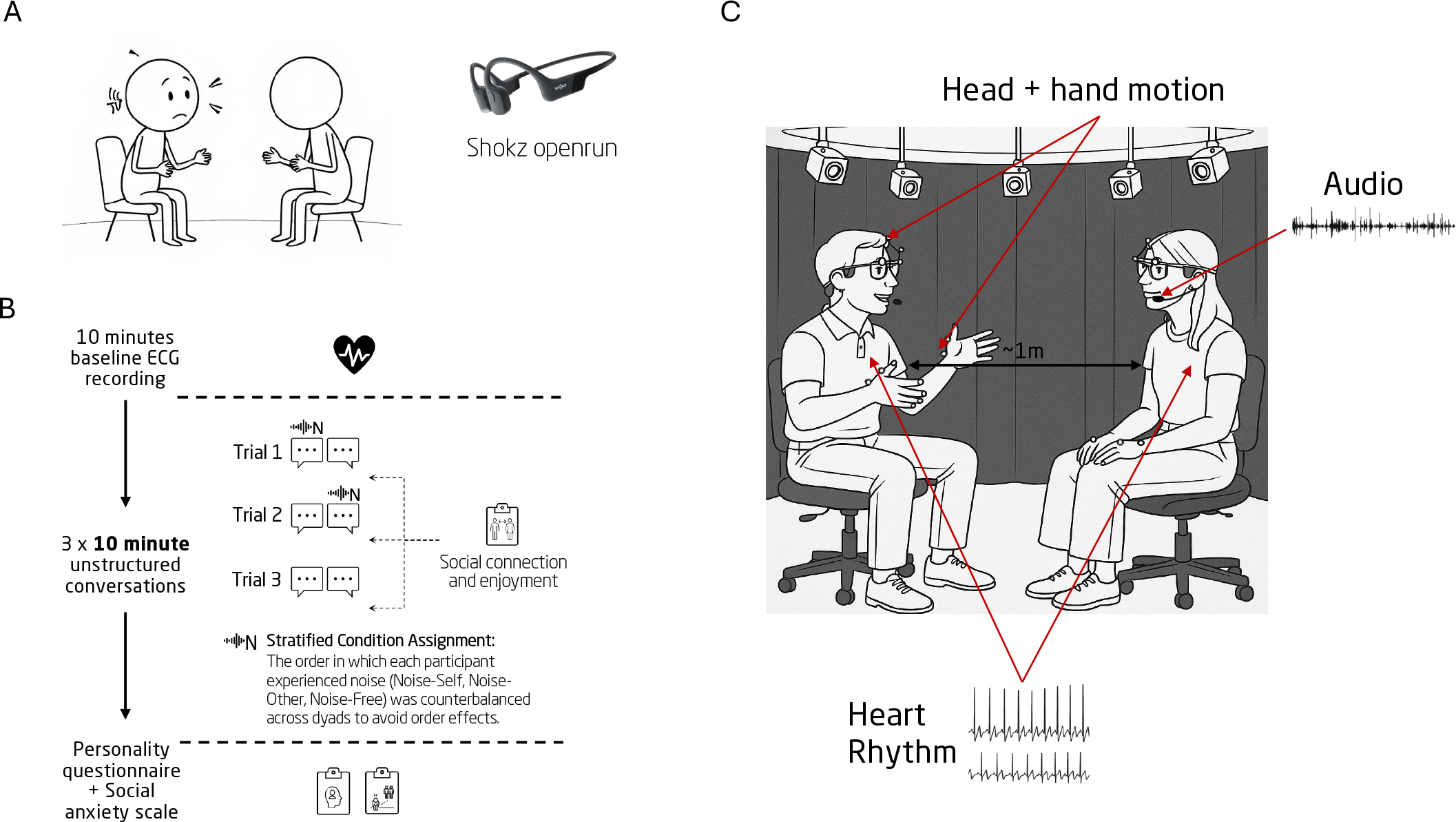
Experimental setup and study design. (A) Asymmetric auditory manipulation using speech-shaped noise delivered through wireless bone conduction headphones, with noise presented to one participant at a time while the other retained normal listening conditions. (B) Experimental procedure. Following a 10-minute resting-state ECG recording, dyads completed three 10-minute unstructured conversations, each followed by a post-conversation questionnaire. Across the three conversations, each participant experienced one Noise-Self, one Noise-Other, and one Noise-Free condition in a counterbalanced order. (C) Experimental setup. Dyads were seated approximately 1 m apart in the conversation booth while speech, head and hand movements, and cardiac activity were recorded simultaneously. *The schematic illustration of the experimental setup was generated using a generative AI tool and is intended for illustrative purposes only*.

The speech-shaped noise was delivered through bone-conduction headphones (Shokz OpenRun). Unlike air-conduction (over-ear or in-ear) headphones, bone-conduction headphones do not occlude the ears, thereby preserving natural auditory cues from the environment. Moreover, they also do not add social distance between interaction partners and remain comfortable during longer conversational tasks. It was used to selectively degrade speech intelligibility through energetic masking while avoiding the additional linguistic and attentional demands associated with competing speech as an interferer, which may alter conversational behaviour beyond perceptual difficulty alone. The noise level was calibrated from masked detection thresholds of the spoken digit “2” as in Weller et al. [35], in an independent sample of 10 adults aged 18–35 years. The speech token was presented at 45 dBA and thresholds were measured using a three-alternative-forced-choice (3-AFC) paradigm with a 1-up, 2-down adaptive procedure, yielding the 70.7% detection point on the psychometric function [52].

Following each conversation, participants completed a short questionnaire assessing their subjective experience of the interaction. At the end of the experiment, participants completed additional questionnaires, including a Big Five personality inventory [53] and the Liebowitz Social Anxiety Scale [54].

### 2.2 Data collection and exclusions

During the experiment, speech, head, and hand movements, as well as cardiac activity were recorded simultaneously. Speech was recorded separately for each participant using head-worn directional microphones (DPA 4188), providing independent audio channels for each speaker. Three-dimensional head and hand movements were captured at 100 Hz using a Vicon motion-capture system (6 Vicon Vero cameras) with reflective 14mm markers attached to the participants’ glasses and hands. Cardiac activity was recorded using Shimmer ECG sensors. Cardiac activity was collected as part of the broader experimental protocol but is not analyzed or reported in the present study. One dyad was excluded because of data-quality issues, leaving 39 dyads and 117 conversations for subjective-rating and turn-taking analyses. Additional trial-level exclusions due to motion-capture errors resulted in 105 trials for interpersonal head coordination, 104 for interpersonal hand coordination, and 110 for intrapersonal hand coordination (see Supplementary S1 for exclusion details).

### 2.3 Data pre-processing and analysis

#### 2.3.1 Subjective ratings

Following each conversation, participants completed a short questionnaire assessing their subjective experience of the interaction. Three measures were collected: perceived conversational flow, enjoyment, and interpersonal closeness. Conversational flow was assessed with the question “How well did this conversation flow?”, and enjoyment was assessed with “How much did you enjoy the conversation you had with your partner?”. Both items were rated on a continuous scale ranging from 0 (“Not at all”) to 100 (“Very” / “Very much”).

Interpersonal closeness was assessed using the Inclusion of Other in the Self (IOS) scale, a single-item measure consisting of seven pairs of circles representing the self and the other, with progressively increasing degrees of overlap between the two circles. Participants selected the pair that best represented their perceived relationship with their conversation partner following each interaction [55].

Responses were analyzed as a function of individual listening condition. Three conditions were defined: *Noise-Self*, in which the participant themselves was exposed to speech-shaped noise; *Noise-Other*, in which the participant’s partner was exposed to noise while the participant remained noise-free; and *Noise-Free*, in which neither participant experienced noise.

#### 2.3.2 Turn-taking

Turn-taking dynamics were extracted from the recorded conversations using a semi-automated annotation procedure combining an automated segmentation and classification pipeline with subsequent human review and correction [56]. A *turn* was defined as a continuous floor-holding contribution by one participant before yielding the conversational floor, whereas a *backchannel* was defined as a listener contribution that provided feedback without attempting to take the floor.

Briefly, speech activity was detected separately for each participant, followed by energy-based cross-talk filtering and temporal merging of speech segments. Candidate conversational units were transcribed using automatic speech recognition and classified as turns or backchannels based on lexical entropy and conversational context [56]. The automatically generated annotations for all 117 conversations were subsequently manually reviewed and corrected before extraction of the turn-taking measures. Further details of the pipeline and annotation procedure are provided in Supplementary S2 and [56].

From the corrected annotations, we extracted mean turn duration, speaker contribution, backchannel frequency, and floor-transfer offsets (FTOs). Speaker contribution was defined as the proportion of the dyad’s total speaking time produced by each participant. FTO was defined as the temporal difference between the offset of one speaker’s turn and the onset of the next speaker’s turn, with positive values indicating gaps and negative values indicating overlap [57]. Backchannels and contributions fully embedded within an interlocutor’s turn were excluded from FTO calculations because they did not represent transitions of the conversational floor.

#### 2.3.3 Movement

##### Pre-processing

Motion capture data were recorded at 100 Hz using a Vicon system. For each participant, three-dimensional trajectories of the head and hands were reconstructed by computing the center of mass (COM) across markers (six markers for the head; two markers per hand). Missing samples were linearly interpolated, signals were low-pass filtered (4th-order Butterworth, 2 Hz cutoff), and time series were downsampled to 10 Hz using Largest-Triangle-Three-Buckets (LTTB) downsampling. Further preprocessing details and data-quality information are provided in Supplementary S4.

##### Multidimensional cross-recurrence quantification analysis (MDCRQA)

Interpersonal and in-trapersonal movement coordination were quantified using multidimensional cross-recurrence quantification analysis (MDCRQA) [58], which captures recurrent states between multivariate movement signals over time. Unlike measures that directly quantify similarity in movement amplitude or velocity, MDCRQA captures whether and when similar states in movement trajectories recur over time, making it less directly driven by overall movement magnitude. Nevertheless, we additionally tested whether the observed recurrence patterns could be accounted for by differences in mean movement speed (Supplementary S5).

For interpersonal head and hand coordination, multivariate time series were constructed from the *x, y*, and *z* trajectories of the respective effectors for each participant. P2 signals were mirrored along the *y*-axis to ensure a common spatial reference frame across partners. In asymmetric noise trials, signals were consistently oriented such that the noise-exposed participant was assigned as the reference time series and the noise-free partner as the second time series.

For intrapersonal hand coordination, left- and right-hand *x, y*, and *z* trajectories were combined to quantify within-person coordination between the two hands. All signals were z-standardized before analysis. MDCRQA parameters were estimated separately for each trial using a data-driven optimization procedure targeting a recurrence rate of 2.9–3.1%. Full parameter-selection and implementation procedures are provided in Supplementary S4.

##### Diagonal cross-recurrence profiles (DCRP)

The temporal structure of interpersonal head and hand coordination was examined using diagonal cross-recurrence profiles (DCRP) [59], which quantify recurrence as a function of temporal lag. For each trial, recurrence was calculated over a lag window from −5 to +5 s. Because the noise-exposed participant was consistently used as the reference signal, negative lags indicate that movements of the noise-exposed participant preceded those of the noise-free partner, whereas positive lags indicate the reverse.

To assess whether observed coordination exceeded chance levels, surrogate DCRP profiles were generated by randomly permuting the temporal order of one participant’s multivariate movement time series while preserving the observed distribution of movement states [60]. The resulting shuffled profiles provide a baseline estimate of recurrence expected in the absence of temporal coupling. Details of the surrogate procedure are provided in Supplementary S4. As participant assignment was arbitrary in the Noise-Free condition, there was no meaningful directional lag orientation. Noise-Free DCRP profiles were therefore symmetrized by averaging each trial’s profile with its time-reversed counterpart.

For intrapersonal hand coordination, DCRPs were calculated between the left- and right-hand movement signals over the same −5 to +5 s lag window. For statistical analysis, recurrence values were averaged within 1-s signed lag bins, retaining temporal information without assuming symmetry around zero lag. Further details are provided in Supplementary S4.

### 2.4 Statistical analysis

All analyses were conducted using mixed-effects models to account for the repeated-measures structure of the data. Unless otherwise specified, linear mixed-effects models were estimated using restricted maximum likelihood (REML), with fixed effects evaluated using Type III ANOVA and Satterthwaite-approximated degrees of freedom. Significant omnibus effects were followed by Tukey-adjusted pairwise comparisons of estimated marginal means (EMMs). Cohen’s *d* was calculated using the residual standard deviation of the corresponding model. Full model specifications and statistical outputs are provided in Supplementary S7.

#### Subjective ratings

Conversational flow and enjoyment were analyzed using linear mixed-effects models with listening condition (*Noise-Self, Noise-Other, Noise-Free*) as a fixed effect and random intercepts for dyad and trial nested within dyad. Interpersonal closeness (IOS) was analyzed using a cumulative link mixed model with trial number, listening condition, and their interaction as fixed effects and a random intercept for dyad. Effects in the ordinal model were evaluated using Type III Wald *χ*^2^ tests.

##### Turn-taking and conversational enjoyment

Average turn duration, speaker contribution, and floor-transfer offset (FTO) were analyzed separately using linear mixed-effects models with individual listening condition (*Noise-Self, Noise-Other, Noise-Free*) as a fixed effect and random intercepts for participant and dyad.

Backchannel counts were analyzed using a negative-binomial mixed-effects model with listening condition as a fixed effect and random intercepts for participant and dyad. The interaction partner’s total speaking time was included as an exposure offset to account for differences in the opportunity to backchannel. The overall effect of condition was evaluated using a Type III Wald *χ*^2^ test, followed by Tukey-adjusted pairwise comparisons of estimated rate ratios.

To examine the relationship between turn-transition timing and conversational enjoyment, valid FTOs were averaged across both participants within each conversation and related to the dyad’s mean enjoyment rating. Enjoyment was modeled as a function of mean FTO, condition (*Noise* vs. *Noise-Free*), and their interaction, with a random intercept for dyad.

##### Interpersonal movement coordination

Interpersonal head and hand DCRP recurrence values were analyzed separately using linear mixed-effects models with condition (*Noise* vs. *Noise-Free*), continuous temporal lag (−5 to +5 s), and their interaction as fixed effects, with random intercepts for dyad and trial. The Condition *×* lag interaction tested whether the temporal structure of coordination differed between listening conditions.

To assess directional asymmetries, lags were additionally grouped into 1-s bins and matching negative and positive lag bins were contrasted within each condition using EMMs with False Discovery Rate (FDR) correction for multiple comparisons.

##### Intrapersonal hand movement coordination

Intrapersonal hand DCRP recurrence was averaged within 1-s signed lag bins across the *±*5-s profile. Recurrence was analyzed using a linear mixed-effects model with listening condition (*Noise-Self, Noise-Other, Noise-Free*) and lag bin as fixed effects. Random intercepts were included for participant, conversation, and participant-specific DCRP profile.

## 3 Results

### 3.1 Asymmetric noise reduced conversational flow and enjoyment

Subjective ratings differed across listening conditions for conversational flow, enjoyment, and perceived interpersonal closeness (Figure 2). For conversational flow, there was a significant main effect of condition (*F* (2, 104.9) = 14.98, *p <* .001). Ratings were lower in both Noise-Other and Noise-Self compared to Noise-Free conditions (*b* = −0.559, *SE* = 0.145, *t* = −3.855, *p* = .0006, *d* = −0.693; *b* = −0.786, *SE* = 0.145, *t* = −5.424, *p <* .001, *d* = −0.975, respectively), while no difference was observed between Noise-Other and Noise-Self (*b* = −0.227, *SE* = 0.129, *t* = −1.762, *p* = .187, *d* = −0.282).

**Figure 2.**
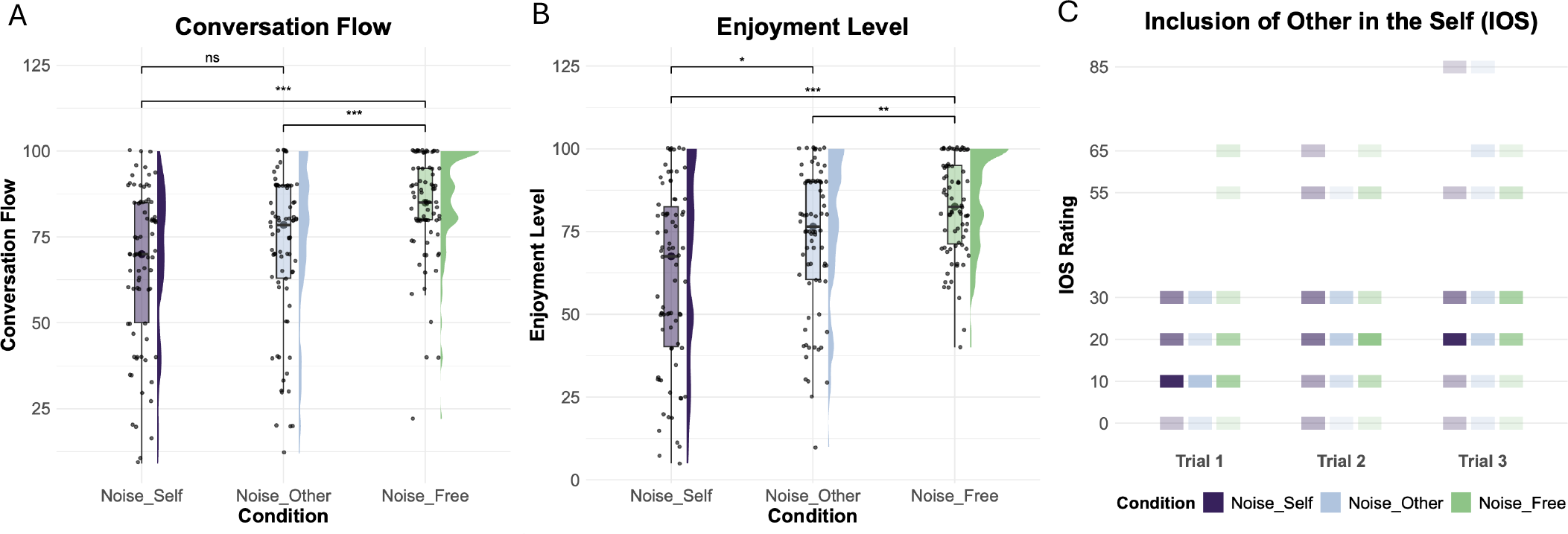
Effect of asymmetric listening conditions on subjective conversational experience. (A) Conversational flow ratings, (B) enjoyment ratings, and (C) interpersonal closeness (IOS) across listening conditions: Noise-Self (participant exposed to noise), Noise-Other (partner exposed to noise), and Noise-Free (neither participant exposed to noise). Both conversational flow and enjoyment were lower under asymmetric listening conditions than in the Noise-Free condition, with the largest reduction in enjoyment observed when participants themselves were exposed to noise. Points in (A) and (B) represent individual participant ratings, boxplots summarize the distribution (median and interquartile range), and significance brackets indicate Tukey-adjusted post hoc pairwise comparisons. For (C), tiles display discrete IOS ratings across trial blocks, with color opacity corresponding to response frequency.

Enjoyment showed a similar effect of condition (*F* (2, 116.65) = 20.17, *p <* .001), with highest ratings in the Noise-Free condition and lower perceived enjoyment in both asymmetric listening conditions. Compared to Noise-Free, enjoyment was lower in both Noise-Self (*b* = −0.907, *SE* = 0.144, *t* = −6.304, *p <* .001, *d* = −1.069) and Noise-Other conditions (*b* = −0.411, *SE* = 0.144, *t* = −2.858, *p* = .014, *d* = −0.485), and also differed between the two noise conditions (*b* = −0.496, *SE* = 0.136, *t* = −3.651, *p* = .001, *d* = −0.585).

Interpersonal closeness, measured via the ordinal Inclusion of Other in the Self (IOS) scale, demonstrated a significant main effect of trial number (*χ*^2^(1) = 21.38, *p <* .001), reflecting increase in perceived closeness over time. However, there was no significant main effect of listening condition (*χ*^2^(2) = 1.95, *p* = .378) nor a condition-by-trial interaction (*χ*^2^(2) = 0.65, *p* = .724), indicating that asymmetric acoustic noise did not modulate the perceived interpersonal closeness over successive trials.

Individual differences in personality did not moderate these effects; linear mixed-effects and ordinal regression models revealed no significant main effects of trait Extraversion nor any Condition × Extraversion interactions on perceived conversational flow, enjoyment, or interpersonal closeness (*ps ≥* .165; Supplementary Figure S3). This confirms that the subjective disruptions induced by asymmetric noise were not driven by individual differences in extroversion.

Overall, while acoustic noise reduced perceived conversational flow and enjoyment, particularly when participants themselves were noise-exposed, interpersonal closeness steadily increased over time regardless of listening condition.

### 3.2 Noise modulated turn-taking dynamics

Turn-taking behaviour differed across listening conditions, revealing systematic asymmetries between noise-exposed participants and their noise-free partners (Figure 3). Average turn duration varied across conditions (*F* (2, 193) = 3.86, *p* = .023). The noise-exposed participant produced longer turns (*M* = 7.20 s) than their noise-free partner (*M* = 6.24 s; *b* = 0.95, *SE* = 0.37, *t* = 2.56, *p* = .030, *d* = 0.41). No significant difference was found between the Noise-Free and the noise-self or noise-other condition (all *p >* .05).

**Figure 3.**
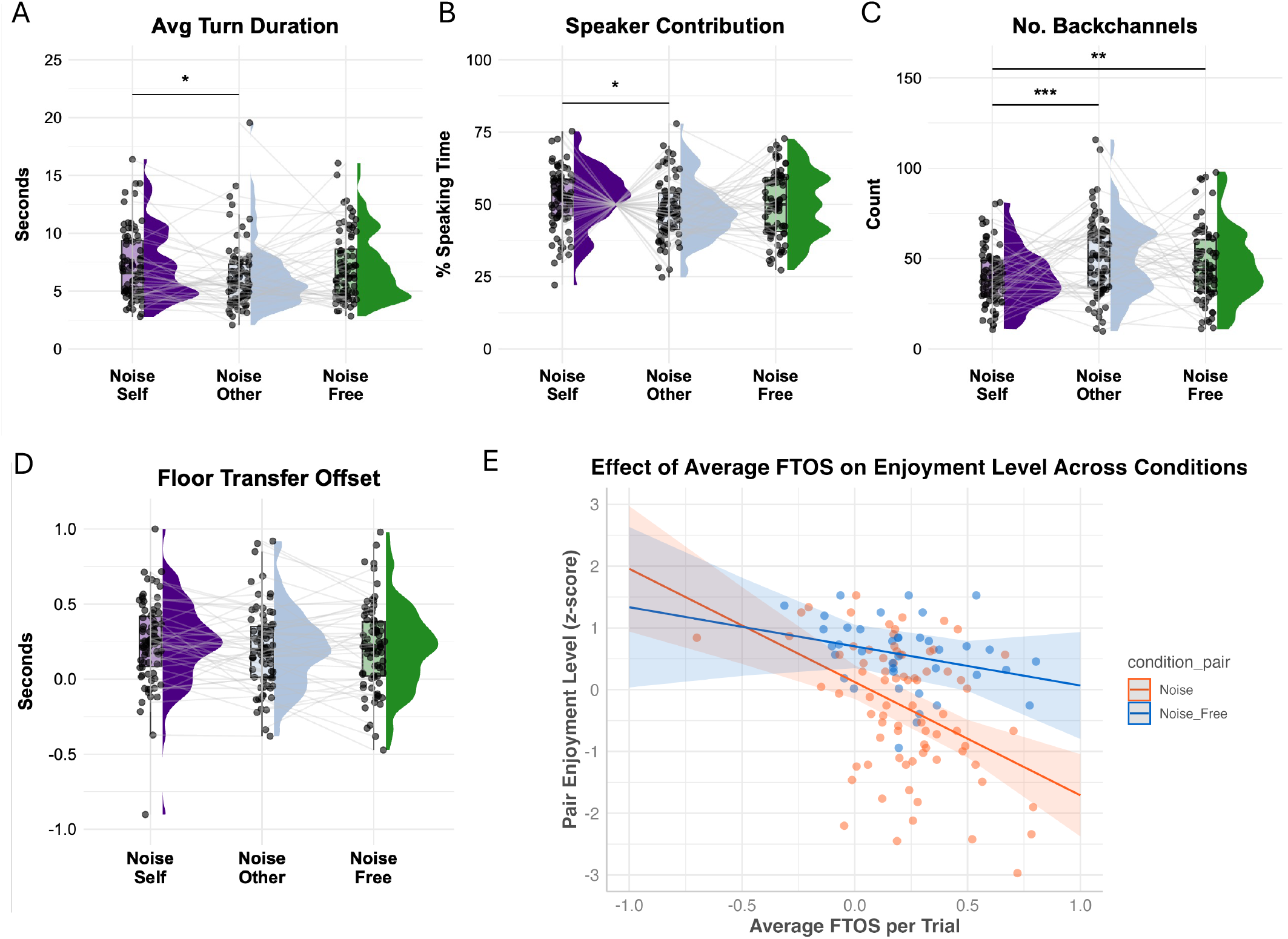
Effects of asymmetric listening conditions on verbal turn-taking behaviour and relation to enjoyment. (A) Average turn duration, (B) speaker contribution (proportion of total speaking time), (C) number of backchannels, and (D) floor-transfer offset (FTO) across listening conditions: Noise-Self (participant exposed to noise), Noise-Other (partner exposed to noise), and Noise-Free (neither participant exposed to noise). Noise-exposed participants took longer turns and spoke for greater proportion per conversation, while their noise-free partners produced more backchannels. Overall FTOs remained comparable across conditions. (E) Association between average floor-transfer offset (FTO) and mean enjoyment ratings across dyads. Shorter FTOs (faster turn transitions) were associated with higher reported enjoyment, with no interaction by listening condition. Points represent individual conversation trials; line indicates model fit with 95% confidence interval.

A similar pattern emerged for speaker contribution (*F* (2, 158.52) = 3.96, *p* = .021). Noise-exposed participants accounted for a larger proportion of total speaking time (*M* = 51.9%) than their noise-free partners (*M* = 48.1%; *b* = 3.88, *SE* = 1.38, *t* = 2.82, *p* = .015, *d* = 0.45). Speaker contribution in the Noise-Free condition (*M* = 50.0%) did not differ significantly from either the Noise-Self (*p* = .340) or Noise-Other condition (*p* = .340).

Backchanneling differed significantly across listening conditions, also after accounting for the interaction partner’s speaking time, *χ*^2^(2) = 25.54, *p <* .001. Relative to the amount of partner speech, noise-exposed participants produced fewer backchannels than their non-exposed partners (rate ratio = 0.86, *z* = −4.97, *p <* .001), corresponding to an approximately 14% reduction. Noise-exposed participants also produced fewer backchannels than participants in the Noise-Free condition (rate ratio = 0.90, *z* = −3.44, *p* = .002). The non-exposed participants did not differ significantly from the Noise-Free condition (rate ratio = 1.05, *z* = 1.53, *p* = .279). An additional analysis directly comparing backchannels per minute of partner speaking time showed the same pattern of condition differences (see Supplementary S6).

In contrast, turn-transition timing showed no significant difference across listening conditions (*F* (2, 193) = 0.95, *p* = .389). Average floor-transfer offset (FTO) remained comparable across Noise-Self (*M* = 0.245 s), Noise-Other (*M* = 0.206 s), and Noise-Free (*M* = 0.210 s) conditions (all post hoc comparisons *p ≥* .425, *d ≤* 0.20).

Together, these results indicate that asymmetric noise perturbed the balance of conversational roles. Noise-exposed participants spoke for a larger proportion of the conversation and produced longer turns, whereas their noise-free partners contributed more frequent listener feedback through verbal backchannelling, suggesting a shift toward more complementary coordination dynamics without altering overall turn-transition latencies.

Furthermore, turn-transition timing was associated with conversational enjoyment (Figure 3E). Across conversations, average floor-transfer offset (FTO) significantly predicted enjoyment ratings, *F* (1, 80.70) = 12.50, *p <* .001. Conversations characterised by faster turn transitions (i.e., shorter FTOs) were generally experienced as more enjoyable than conversations with longer delays between speakers. Crucially, there was no significant interaction between FTO and listening condition on enjoyment, indicating that this positive relationship between rapid turn-taking and conversational enjoyment remained consistent across both noise-free and asymmetric noise conditions. This finding is consistent with previous work showing that faster response timing was related to greater feelings of social connection and conversational quality [16].

Analysis of personality traits revealed that while higher trait Extraversion was associated with greater overall speaking contribution (*F* (1, 219.00) = 5.84, *p* = .017), it did not significantly predict turn duration (*F* (1, 217.90) = 1.08, *p* = .299), backchannel frequency (*F* (1, 214.13) = 1.15, *p* = .286), or floor-transfer offset (*F* (1, 209.90) = 2.91, *p* = .090). Moreover, no significant Extraversion *×* Condition interactions were observed across any turn-taking features (*ps ≥* .216; Supplementary Figure S4).

### 3.3 Interpersonal head movement coordination showed leader-follower like dynamics under asymmetric noise

When comparing interpersonal movement coordination strength across conditions, no significant main effect of condition was observed (*F* = 1.09, *p* = .30), indicating that overall recurrence strength did not differ between Noise and Noise-Free conditions. A significant main effect of lag was observed (*F* = 55.14, *p <* .001), reflecting systematic variation in coordination strength across temporal delays.

Importantly, a significant Condition *×* lag interaction was found (*F* = 55.14, *p <* .001), indicating that the temporal structure of coordination differed between conditions (Figure 4A). This interaction remained unchanged when controlling for head movement magnitude, which did not significantly differ between conditions (*p* = .161) or predict recurrence (*p* = .317; Supplementary S5). As illustrated in Figure 4A, the noise-free condition showed balanced recurrence across corresponding negative and positive lags, with peaks at approximately −3 s and +3 s. Post hoc symmetry contrasts demonstrated that recurrence in the noise-free condition was balanced across all matching negative and positive lag windows (all |*b*| *≤* 1.03 *×* 10^™4^, |*z*| *≤* 0.30, *p*_adj_ = .962). This pattern indicates a bidirectional coordination structure, in which coupling strength is distributed symmetrically across temporal delays.

**Figure 4.**
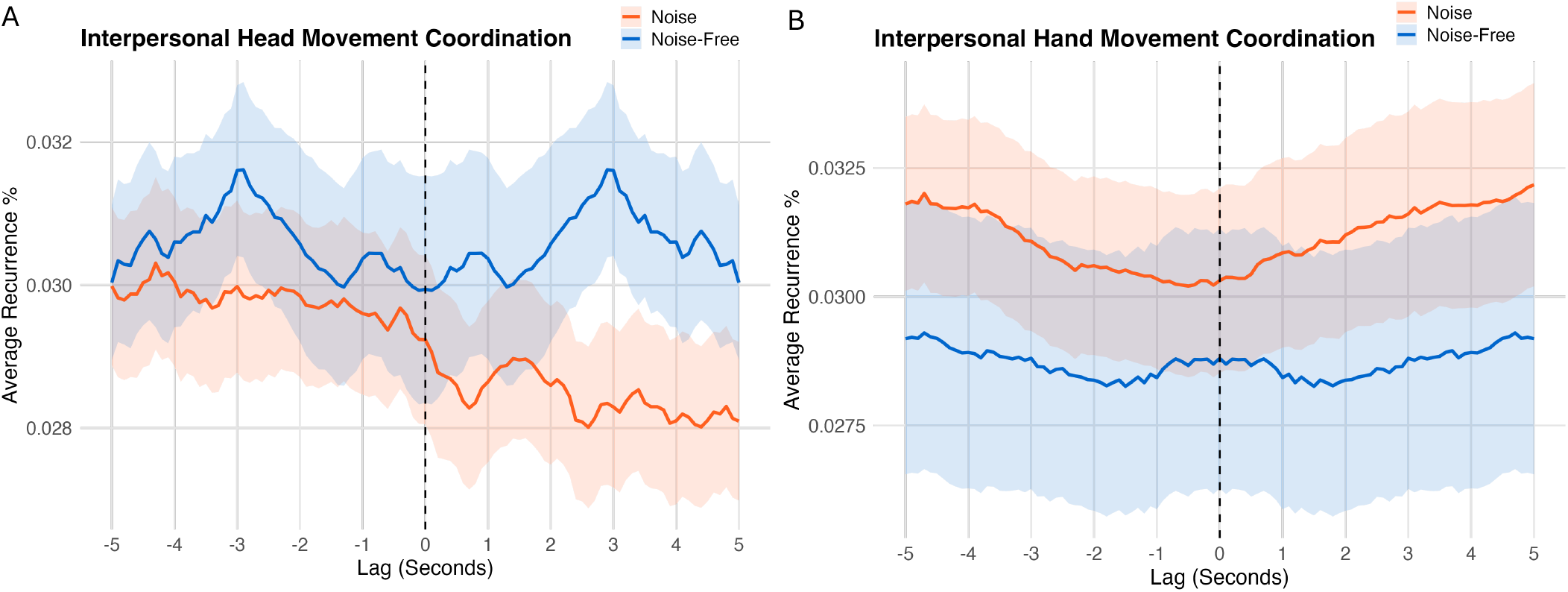
Interpersonal movement coordination across conditions. (A) Diagonal cross-recurrence profiles (DCRP) of head movements across temporal lags in Noise and Noise-Free conditions. The Noise-Free condition shows a relatively symmetric lag structure, whereas the asymmetric Noise condition shows higher recurrence at negative lags, indicating that movements of the noise-exposed participant more often preceded those of their partner. (B) DCRP of interpersonal hand movements across conditions. Hand-movement recurrence was low and relatively flat across lags, with no significant differences in overall recurrence or lag-dependent structure between conditions. Shaded areas indicate standard error.

In comparison, the asymmetric noise condition displayed a significantly asymmetric, monotonic lag profile, characterised by higher recurrence in the negative relative to the positive lag region. Post hoc symmetry contrasts revealed significantly greater recurrence at negative delays compared to their corresponding positive counterparts across the entire lag window (e.g., [−3 to − 2 s] − [2 to 3 s]: *b* = 1.57 *×* 10^™3^, *SE* = 2.30 *×* 10^™4^, *z* = 6.84, *p*_adj_ *<* .0001; [−4 to − 3 s] − [3 to 4 s]: *b* = 1.54 *×* 10^™3^, *SE* = 2.30 *×* 10^™4^, *z* = 6.70, *p*_adj_ *<* .0001; all *p*_adj_ *≤* .0002). Given that negative lag values correspond to instances in which the head movements of the noise-exposed participant precede those of their partner, this statistical asymmetry indicates that the noise-exposed participant led the temporal dynamics of the interaction.

For hand movements, there was no significant main effect of condition (*F* (1, 102) = 0.63, *p* = .430), indicating that overall interpersonal hand recurrence did not differ between asymmetric-noise and noise-free conversations. There was also no significant main effect of lag (*F* (1, 10398) = 2.74, *p* = .098) and no significant Condition *×* lag interaction (*F* (1, 10398) = 2.74, *p* = .098). As shown in Figure 4B, the DCRP profiles remained relatively flat across temporal lags in both conditions, with no clear lag-dependent peaks or asymmetries.

Notably, hand movement magnitude was greater in the Noise condition than in the Noise-Free condition (*p* = .035). However, controlling for hand movement magnitude did not alter the recurrence results, and hand movement magnitude was not significantly associated with recurrence (*p* = .451; Supplementary S5). Taken together, these results indicate that, unlike head movements, the temporal structure of interpersonal hand coordination did not significantly differ between conditions, despite greater hand movement magnitude during Noise conversations.

### 3.4 Intrapersonal hand movement coordination

As illustrated in Figure 5, intrapersonal hand recurrence varied across temporal lags and listening conditions. Recurrence was highest overall in the *Noise-Self* condition, whereas the *Noise-Other* and *Noise-Free* profiles were more similar.

**Figure 5.**
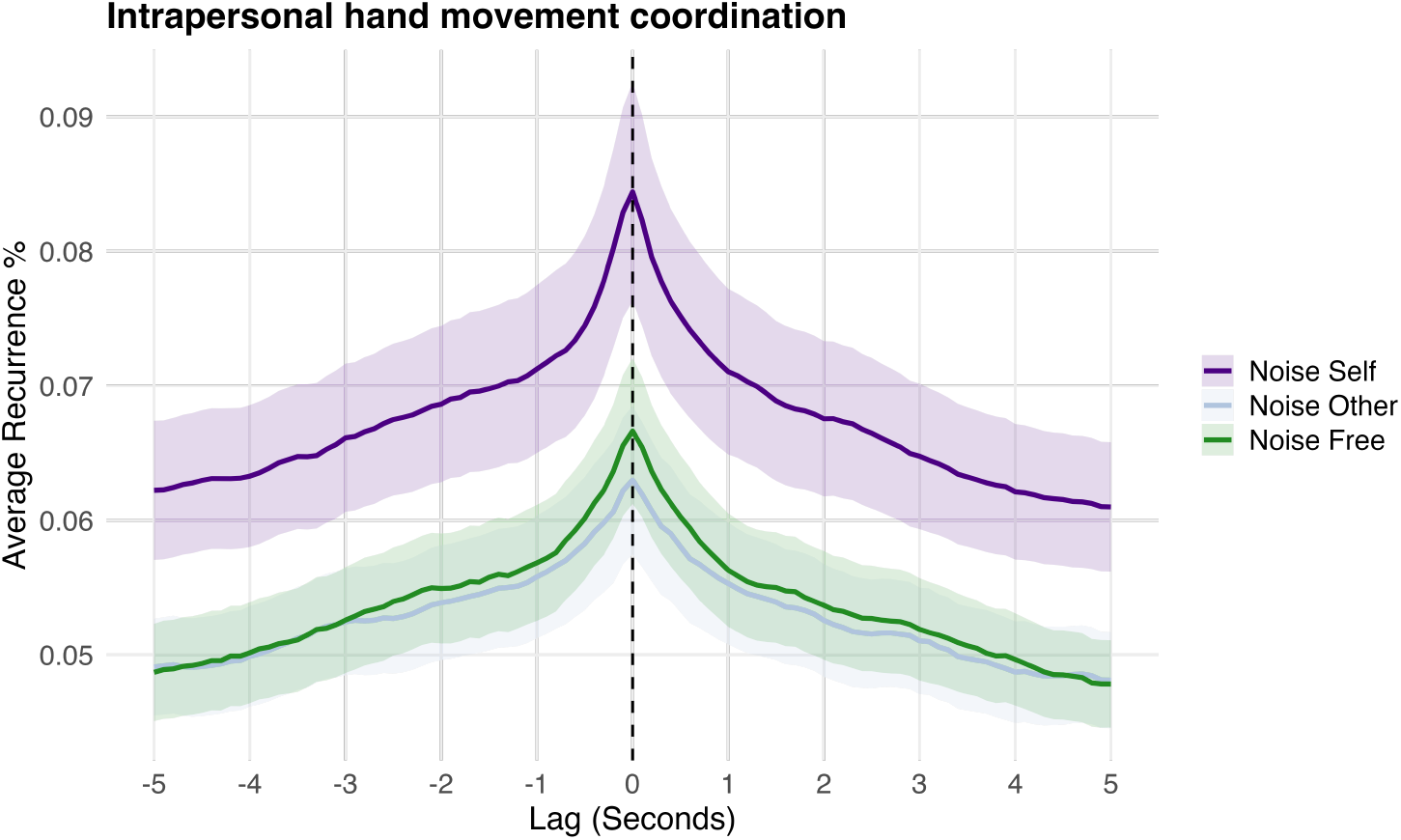
Intrapersonal hand movement coordination across conditions. Diagonal recurrence profiles (DCRP) of bilateral hand movements within individuals in the Noise-Self, Noise-Other, and Noise-Free conditions. Recurrence is shown across signed temporal lags from −5 to +5 s. Shaded areas indicate standard error.

A linear mixed-effects model revealed a significant main effect of listening condition on intrapersonal recurrence (*F* (2, 134.91) = 3.68, *p* = .028). Recurrence also differed significantly across lag bins (*F* (9, 1836.00) = 57.04, *p <* .001), indicating clear temporal structure in the DCRP profiles.

Estimated marginal means were highest in the *Noise-Self* condition (*M* = 0.0677, *SE* = 0.0046), followed by *Noise-Free* (*M* = 0.0542, *SE* = 0.0046) and *Noise-Other* (*M* = 0.0533, *SE* = 0.0046). Tukey-adjusted pairwise comparisons showed higher recurrence in *Noise-Self* than *Noise-Other* (*β* = 0.0144, *p* = .048). The difference between *Noise-Self* and *Noise-Free* was only marginally significant after correction (*β* = 0.0135, *p* = .064), and *Noise-Other* did not differ from *Noise-Free* (*β* = −0.0009, *p* = .988).

These results indicate that intrapersonal hand coordination varied with the participant’s role under asymmetric listening conditions. In particular, recurrence was higher when participants themselves experienced noise than when their partner experienced noise, although the difference relative to the Noise-Free baseline was not statistically significant after correction. Furthermore, this pattern remained unchanged when controlling for participants’ hand movement magnitude (Supplementary S5).

## 4 Discussion

The present study investigated how asymmetric auditory constraints influence coordination during natural conversation. Across subjective, verbal, and movement measures, asymmetric noise systematically modulated conversational dynamics. Participants exposed to noise reported poorer flow and enjoyment, produced longer turns, and contributed more speaking time, while their noise-free partners produced more backchannels. Interpersonal head coordination became temporally asymmetric, with movements of the noise-exposed participant more often preceding those of their partner. Interpersonal hand recurrence did not show a significant overall difference between conditions nor any change in temporal structure, and intrapersonal hand coordination was stronger when participants themselves experienced noise than when their partner experienced noise. Together, these findings show that unequal listening conditions lead conversation partners to adjust in different ways across verbal and nonverbal channels.

One of the clearest effects was a shift in verbal participation. Participants experiencing degraded auditory input held the floor for longer and contributed more speaking time, whereas their partners produced more backchannels. Listening under degraded conditions increases cognitive load and may make it harder to monitor incoming speech while planning responses [43, 61]. This may make speakers slower or less likely to hand over the conversational floor, leading to longer speaking turns [12, 62]. Conversely, the noise-free partner may provide more listener feedback to support grounding. This pattern is consistent with joint-action research showing that unequal information or task constraints can produce complementary rather than symmetric adaptation [40, 63, 64].

The movement analyses showed that the effects of asymmetric listening differed across movement modalities. Head coordination became temporally asymmetric under noise, in contrast to the more symmetric coupling observed during noise-free conversation. Although earlier movement timing by one partner is not equivalent to leadership, it indicates reduced symmetry in how partners adapted their head movements to one another. Hand coordination showed no corresponding effect where overall recurrence did not differ between listening conditions, and the temporal profiles showed no clear lag-dependent structure. This difference may reflect distinct communicative functions of head and hand movements. Head movements are closely linked to turn regulation, listener feedback, and conversational timing [9, 18, 19, 26], whereas hand movements more often convey semantic and pragmatic information alongside speech [65, 66]. Unequal listening conditions may therefore influence the timing of head coordination more strongly than interpersonal hand coordination.

Interestingly, hand movement magnitude increased under noise, possibly suggesting a shift to non-verbal communication.

Unequal listening conditions also affected coordination within individuals. Participants exposed to noise showed stronger coordination between their own hands than when their partner experienced noise. The present data cannot determine the underlying mechanism, but this pattern may reflect greater self-oriented motor organisation under increased perceptual demands. This interpretation is consistent with recent work showing that interpersonal interaction can influence intrapersonal coordination. Intrapersonal bodily rhythms can become more out of phase with each other when people synchronize interpersonally with another person’s rhythms [48], suggesting that social interaction can influence how coordination is organised within the individual. This interpretation remains speculative, but it could be examined further by combining interpersonal and intrapersonal movement measures with physiological or neurocognitive indices of listening effort.

Subjective ratings indicate that these behavioural adjustments did not fully offset the consequences of degraded audio. Participants reported poorer flow and enjoyment during asymmetric noise, particularly when they themselves experienced the manipulation. Faster floor-transfer offsets were also associated with greater enjoyment, replicating previous work linking response timing to conversational quality and social affiliation [16, 17]. This relationship did not differ across listening conditions, suggesting that turn-transition timing remains strongly linked to conversational experience even when listening conditions are unequal.

Several limitations should be acknowledged. First, participants were young adults with normal hearing interacting with unfamiliar partners. Different adaptation strategies may emerge in older adults, individuals with hearing impairment, or established relationships in which conversational routines are already well developed. Second, the manipulation consisted of experimentally controlled speech-shaped noise during relatively short laboratory conversations. Although this allowed precise manipulation of listening difficulty while minimizing confounds associated with competing speech, real-world listening environments are more variable and may involve fluctuating noise, reverberation, competing speakers, or chronic hearing difficulties [67].

A further limitation is the absence of a condition in which both participants experienced noise. While our design allowed us to examine role-specific adaptation within asymmetric interactions, comparisons with the Noise-free condition cannot distinguish effects of perceptual asymmetry from those of listening difficulty more generally. Future studies incorporating a shared-noise condition could help disentangle these effects.

The use of unstructured conversation also presents an important trade-off. Allowing participants to converse freely provided a naturalistic setting in which coordination could emerge spontaneously, but introduced substantial variability in conversational content and behaviour across trials. More structured conversational tasks could provide greater experimental control and help isolate specific coordination mechanisms, although constraining the interaction may also engage different coordination processes than those emerging during free conversation.

The movement analyses likewise provide a relatively coarse-grained characterisation of coordination. MDCRQA has the advantage of preserving multidimensional movement information without reducing behaviour to predefined movement events, but recurrence between high-dimensional spatial trajectories does not directly identify the communicative function of particular movements. Moreover, modelling continuous recurrence profiles across both condition and temporal lag remains statistically challenging, and the present analyses may not capture all changes in the shape and timing of coordination. Approaches that combine recurrence analysis with movement-event classification, conversation unit annotation, or complementary time-series models may provide a more detailed account of how coordination changes across behavioural channels.

Finally, although the directional head-movement patterns indicate asymmetric temporal organisation, they cannot establish causal influence between partners. Future work using methods designed to quantify directional influence could more directly test how adaptation between interlocutors changes under asymmetric listening conditions.

Nevertheless, a key strength of our study is the use of a controlled perturbation during natural, unstructured conversation, allowing us to investigate how coordination dynamics change under unequal interaction demands. By combining verbal and nonverbal measures, and examining coordination both within and between individuals, we could capture how different aspects of conversational coordination are reorganised in response to the perturbation.

Taken together, our findings extend previous work on nonverbal joint action to natural conversation, showing that unequal interaction demands can lead to complementary patterns of coordination between partners. Importantly, these changes were not uniform across modalities: verbal contributions and the temporal structure of head-movement coordination became asymmetric, while turn-transition timing and interpersonal hand-movement coordination remained largely unchanged. Rather than testing whether coordination increases or decreases under communicative difficulty, these findings highlight the importance of examining who adapts, how, and through which behavioural channels. Coordination is therefore better understood as a dynamic process in which partners adjust their contributions in relation to one another across modalities, which becomes reorganised within and between individuals when interaction demands are unequal.

## Data accessibility

The data and analysis code supporting the findings of this study are available here: https://figshare.com/s/b60a377b4293f5c6abfb

## Ethics

All participants provided informed consent for participation and for audio, video, ECG, and motion recording. The study was approved by the Science-Ethics Committee for the Capital Region of Denmark (reference H-16036391) and conducted in accordance with the Declaration of Helsinki. Participants additionally provided written consent for the public sharing of pseudo-anonymised data via DTU’s data repository, including ECG, head and hand movement data, and questionnaire responses. They were informed that these data would be fully de-identified and not linked to personal information, and that audio and video recordings would not be shared. Participants were informed of GDPR-related data processing and their rights under EU regulation 2016/679, including the right to access, correct, and withdraw their data.

## Authors’ contributions

H.H.: Conceptualization, Methodology, Software, Formal analysis, Investigation, Data curation, Writing – original draft, Writing – review & editing. A.A.: Conceptualization, Methodology, Investigation, Resources, Writing – review & editing. M.M.: Methodology, Investigation, Resources. R.T.: Methodology, Formal analysis. I.K.: Conceptualization, Methodology, Supervision, Funding acquisition, Writing – review & editing.

## Declaration of AI use

Generative AI tools were used for language editing and refinement of the manuscript and to generate the schematic experimental illustration in Figure 1. All scientific content, analyses, interpretations, and conclusions were reviewed and verified by the authors.

## References

[1] Herbert H Clark. Using language. Cambridge university press, 1996.

[2] Lauren V Hadley, Graham Naylor, and Antonia F de C Hamilton. A review of theories and methods in the science of face-to-face social interaction. Nature Reviews Psychology, 1(1):42–54, 2022.

[3] Natalie Sebanz, Harold Bekkering, and Günther Knoblich. Joint action: bodies and minds moving together. Trends in cognitive sciences, 10(2):70–76, 2006.

[4] Simon Garrod and Martin J Pickering. Joint action, interactive alignment, and dialog. Topics in Cognitive Science, 1(2):292–304, 2009.

[5] Susan E Brennan, Alexia Galati, and Anna K Kuhlen. 8 two minds, one dialog: Coordinating speaking and understanding. Psychology of learning and motivation, 53:301, 2010.

[6] Max M Louwerse, Rick Dale, Ellen G Bard, and Patrick Jeuniaux. Behavior matching in multimodal communication is synchronized. Cognitive science, 36(8):1404–1426, 2012.

[7] Kevin Shockley, Daniel C Richardson, and Rick Dale. Conversation and coordinative structures. Topics in Cognitive Science, 1(2):305–319, 2009.

[8] Alexandra Paxton, Rick Dale, and Daniel C Richardson. 18 social coordination of verbal and nonverbal behaviours. Interpersonal coordination and performance in social systems, page 259, 2016.

[9] Judith Holler and Anna K Kuhlen. Psycholinguistic perspectives on face-to-face conversation. Nature Reviews Psychology, pages 1–16, 2026.

[10] Rick Dale, Riccardo Fusaroli, Nicholas D Duran, and Daniel C Richardson. The self-organization of human interaction. In Psychology of learning and motivation, volume 59, pages 43–95. Elsevier, 2013.

[11] Daniel Richardson, Rick Dale, and Kevin Shockley. Synchrony and swing in conversation: Coordination, temporal dynamics, and communication. Embodied communication in humans and machines, pages 75–94, 2008.

[12] Stephen C Levinson. Turn-taking in human communication–origins and implications for language processing. Trends in cognitive sciences, 20(1):6–14, 2016.

[13] Antje S Meyer. Timing in conversation. Journal of Cognition, 6(1):20, 2023.

[14] Carina Riest, Annett B Jorschick, and Jan P de Ruiter. Anticipation in turn-taking: mechanisms and information sources. Frontiers in psychology, 6:89, 2015.

[15] Ivana Konvalinka, Kyveli Kompatsiari, and Qianliang Li. The fine-grained temporal dynamics of social timing: a window into sociality of embodied social agents comment on “the evolution of social timing” by l. verga, sa kotz, & a. ravignani. Physics of Life Reviews, 47:95–98, 2023.

[16] Emma M Templeton, Luke J Chang, Elizabeth A Reynolds, Marie D Cone LeBeaumont, and Thalia Wheatley. Fast response times signal social connection in conversation. Proceedings of the National Academy of Sciences, 119(4):e2116915119, 2022.

[17] Emma M Templeton, Luke J Chang, Elizabeth A Reynolds, Marie D Cone LeBeaumont, and Thalia Wheatley. Long gaps between turns are awkward for strangers but not for friends. Philosophical Transactions of the Royal Society B: Biological Sciences, 378(1875), 2023.

[18] Lorenza Mondada. Multiple temporalities of language and body in interaction: Challenges for transcribing multimodality. Research on language and social interaction, 51(1):85–106, 2018.

[19] Tifenn Fauviaux, Ludovic Marin, Mathilde Parisi, Richard Schmidt, and Ghilès Mostafaoui. From unimodal to multimodal dynamics of verbal and nonverbal cues during unstructured conversation. PLoS One, 19(9):e0309831, 2024.

[20] Alexandra Paxton and Rick Dale. Interpersonal movement synchrony responds to high-and low-level conversational constraints. Frontiers in psychology, 8:1135, 2017.

[21] Shay Ohayon and Ilanit Gordon. Multimodal interpersonal synchrony: Systematic review and meta-analysis. Behavioural Brain Research, 480:115369, 2025.

[22] Judith Holler and Stephen C Levinson. Multimodal language processing in human communication. Trends in cognitive sciences, 23(8):639–652, 2019.

[23] Janet Beavin Bavelas, Linda Coates, and Trudy Johnson. Listener responses as a collaborative process: The role of gaze. Journal of communication, 52(3):566–580, 2002.

[24] Judith Holler, Kobin H Kendrick, and Stephen C Levinson. Processing language in face-to-face conversation: Questions with gestures get faster responses. Psychonomic bulletin & review, 25(5):1900–1908, 2018.

[25] Kobin H Kendrick, Judith Holler, and Stephen C Levinson. Turn-taking in human face-to-face interaction is multimodal: gaze direction and manual gestures aid the coordination of turn transitions. Philosophical Transactions of the Royal Society B: Biological Sciences, 378(1875):20210473, 2023.

[26] Anna K Kuhlen, Alexia Galati, and Susan E Brennano. Gesturing integrates top-down and bottom-up information: Joint effects of speakers’ expectations and addressees’ feedback. Language and Cognition, 4(1):17–41, 2012.

[27] Amélie Rochet-Capellan and Susanne Fuchs. Take a breath and take the turn: how breathing meets turns in spontaneous dialogue. Philosophical Transactions of the Royal Society B: Biological Sciences, 369(1658), 2014.

[28] Christian Rominger, Ilona Papousek, Andreas Fink, Corinna M Perchtold, Helmut K Lackner, Elisabeth M Weiss, and Andreas R Schwerdtfeger. Creative challenge: Regular exercising moderates the association between task-related heart rate variability changes and individual differences in originality. PLoS One, 14(7):e0220205, 2019.

[29] Joana Coutinho, Alfredo Pereira, Patrícia Oliveira-Silva, Deborah Meier, Vladimiro Lourenço, and Wolfgang Tschacher. When our hearts beat together: Cardiac synchrony as an entry point to understand dyadic co-regulation in couples. Psychophysiology, 58(3):e13739, 2021.

[30] Richard V Palumbo, Marisa E Marraccini, Lisa L Weyandt, Oliver Wilder-Smith, Heather A McGee, Siwei Liu, and Matthew S Goodwin. Interpersonal autonomic physiology: A systematic review of the literature. Personality and social psychology review, 21(2):99–141, 2017.

[31] Riccardo Fusaroli, Joanna Rączaszek-Leonardi, and Kristian Tylén. Dialog as interpersonal synergy. New Ideas in Psychology, 32:147–157, 2014.

[32] Denise Haunani Solomon, Miriam Brinberg, Graham Bodie, Susanne Jones, and Nilam Ram. A dynamic dyadic systems perspective on interpersonal conversation. Communication Methods and Measures, 17(4):273–292, 2023.

[33] Marlou Rasenberg, Wim Pouw, Asli Özyürek, and Mark Dingemanse. The multimodal nature of communicative efficiency in social interaction. Scientific Reports, 12(1):19111, 2022.

[34] Rick Dale, Gregory A Bryant, Joseph H Manson, and Matthew M Gervais. Body synchrony in triadic interaction. Royal Society open science, 7(9):200095, 2020.

[35] Tobias Weller, Jörg M Buchholz, and Virginia Best. Auditory masking of speech in reverberant multi-talker environments. The Journal of the Acoustical Society of America, 139(3):1303–1313, 2016.

[36] Kelly Miles, Adam Weisser, Rachel W Kallen, Manuel Varlet, Michael J Richardson, and Joerg M Buchholz. Behavioral dynamics of conversation,(mis) communication and coordination in noisy environments. Scientific reports, 13(1):20271, 2023.

[37] A Josefine Munch Sørensen, Michal Fereczkowski, and Ewen N MacDonald. Effects of noise and second language on conversational dynamics in task dialogue. Trends in Hearing, 25:23312165211024482, 2021.

[38] James Trujillo, Asli Özyürek, Judith Holler, and Linda Drijvers. Speakers exhibit a multimodal lombard effect in noise. Scientific reports, 11(1):16721, 2021.

[39] Ingvi Örnólfsson, Axel Ahrens, Tobias May, and Torsten Dau. Investigating the impact of background noise on collaborative decision-making using an individual-weighted voting model. Cognitive Research: Principles and Implications, 11(1):17, 2026.

[40] Joshua C. Skewes, Lea Skewes, John Michael, and Ivana Konvalinka. Synchronised and complementary coordination mechanisms in an asymmetric joint aiming task. Experimental Brain Research, 233(2):551–565, 2015.

[41] Cordula Vesper, Robrecht PRD Van Der Wel, Günther Knoblich, and Natalie Sebanz. Making oneself predictable: Reduced temporal variability facilitates joint action coordination. Experimental brain research, 211(3):517–530, 2011.

[42] Tehran J Davis, Gabriela B Pinto, and Adam W Kiefer. The stance leads the dance: the emergence of role in a joint supra-postural task. Frontiers in Psychology, 8:718, 2017.

[43] Jonathan E Peelle. Listening effort: How the cognitive consequences of acoustic challenge are reflected in brain and behavior. Ear and hearing, 39(2):204, 2018.

[44] Merle T Fairhurst, Petr Janata, and Peter E Keller. Leading the follower: an fmri investigation of dynamic cooperativity and leader–follower strategies in synchronization with an adaptive virtual partner. Neuroimage, 84:688–697, 2014.

[45] Henning Holle, Jonas Obleser, Shirley-Ann Rueschemeyer, and Thomas C Gunter. Integration of iconic gestures and speech in left superior temporal areas boosts speech comprehension under adverse listening conditions. NeuroImage, 49(1):875–884, 2010.

[46] Christian Obermeier, Thomas Dolk, and Thomas C Gunter. The benefit of gestures during communication: Evidence from hearing and hearing-impaired individuals. Cortex, 48(7):857–870, 2012.

[47] Lauren V Hadley and Jamie A Ward. Synchrony as a measure of conversation difficulty: Movement coherence increases with background noise level and complexity in dyads and triads. PLoS One, 16(10):e0258247, 2021.

[48] Ivana Konvalinka, Natalie Sebanz, and Günther Knoblich. The social, decoupled self: interpersonal synchronization of breathing alters intrapersonal cardiorespiratory coupling. Annals of the New York Academy of Sciences, 1559(1):e70285, 2026.

[49] Laura Galbusera, Michael TM Finn, Wolfgang Tschacher, and Miriam Kyselo. Interpersonal synchrony feels good but impedes self-regulation of affect. Scientific reports, 9(1):14691, 2019.

[50] Ole Adrian Heggli, Joana Cabral, Ivana Konvalinka, Peter Vuust, and Morten L Kringelbach. A kuramoto model of self-other integration across interpersonal synchronization strategies. PLoS computational biology, 15(10):e1007422, 2019.

[51] Camilla Gregorini, Mario Medoni, Anna Zamm, Marco Petilli, Arianna Schiano Lomoriello, and Ivana Konvalinka. Alpha and beta event-related synchronization as neural markers of self-other processing during sensorimotor coordination. Acta Psychologica, 270:107679, 2026.

[52] HCCH Levitt. Transformed up-down methods in psychoacoustics. The Journal of the Acoustical society of America, 49(2B):467–477, 1971.

[53] Boele De Raad. The big five personality factors: the psycholexical approach to personality. Hogrefe & Huber Publishers, 2000.

[54] Michael R Liebowitz. Liebowitz social anxiety scale. Journal of Anxiety Disorders, 1987.

[55] Arthur Aron, Elaine N Aron, and Danny Smollan. Inclusion of other in the self scale and the structure of interpersonal closeness. Journal of personality and social psychology, 63(4):596, 1992.

[56] Hanlu He, Harald Vilhelm Skat-Rørdam, Ingvi Örnólfsson, and Ivana Konvalinka. An automated pipeline for standardised speech-unit annotation in spontaneous dialogue, 2026.

[57] Jan-Peter De Ruiter, Holger Mitterer, and Nick J Enfield. Projecting the end of a speaker’s turn: A cognitive cornerstone of conversation. Language, 82(3):515–535, 2006.

[58] Sebastian Wallot. Multidimensional cross-recurrence quantification analysis (mdcrqa)–a method for quantifying correlation between multivariate time-series. Multivariate behavioral research, 54(2):173–191, 2019.

[59] Sebastian Wallot and Giuseppe Leonardi. Analyzing multivariate dynamics using cross-recurrence quantification analysis (crqa), diagonal-cross-recurrence profiles (dcrp), and multidimensional recurrence quantification analysis (mdrqa)–a tutorial in r. Frontiers in psychology, 9:2232, 2018.

[60] Daniel C Richardson and Rick Dale. Looking to understand: The coupling between speakers’ and listeners’ eye movements and its relationship to discourse comprehension. Cognitive science, 29(6):1045–1060, 2005.

[61] Ruth E Corps, Muzna Shehzad, Andrew I McLaren, Leigh B Fernandez, and Lauren V Hadley. Effects of listening demand on turn-taking processes: Investigating turn-end prediction and verbal response planning. Acta Psychologica, 260:105694, 2025.

[62] A Josefine Munch Sørensen, Thomas Lunner, and Ewen N MacDonald. Conversational dynamics in task dialogue between interlocutors with and without hearing impairment. Trends in Hearing, 28:23312165241296073, 2024.

[63] Cordula Vesper and Michael J Richardson. Strategic communication and behavioral coupling in asymmetric joint action. Experimental brain research, 232(9):2945–2956, 2014.

[64] Michael J Richardson, Steven J Harrison, Rachel W Kallen, Ashley Walton, Brian A Eiler, Elliot Saltzman, and RC Schmidt. Self-organized complementary joint action: Behavioral dynamics of an interpersonal collision-avoidance task. Journal of Experimental Psychology: Human Perception and Performance, 41(3):665, 2015.

[65] Elena T Levy and David McNeill. Speech, gesture, and discourse. Discourse processes, 15(3):277–301, 1992.

[66] Adam Kendon. Gesture: Visible action as utterance. Cambridge University Press, 2004.

[67] M Kathleen Pichora-Fuller, Sophia E Kramer, Mark A Eckert, Brent Edwards, Benjamin WY Hornsby, Larry E Humes, Ulrike Lemke, Thomas Lunner, Mohan Matthen, Carol L Mackersie, et al. Hearing impairment and cognitive energy: The framework for understanding effortful listening (fuel). Ear and hearing, 37:5S–27S, 2016.

