## Supplementary information for "Asymmetric listening conditions perturb verbal and nonverbal coordination dynamics during conversation"

This PDF file includes:

- Supplementary Notes S1–S7
- Supplementary Figures S1–S10
- Supplementary Tables S1–S3
- Supplementary references

**S1. Data retention and exclusions.** A total of 40 dyads (80 participants) completed the experiment, each contributing three conversation trials. One dyad was excluded from all analyses due to data quality issues, resulting in a final sample of 39 dyads and 117 conversation trials.

All 117 trials were retained for the subjective-rating and turn-taking analyses. Movement analyses required additional trial-level exclusions due to motion-capture data quality issues, including calibration and marker reconstruction errors. Exclusions were applied separately for each movement analysis because the availability and quality of head and hand marker trajectories differed across trials.

For interpersonal head movement coordination, 105 of 117 trials were retained (72 asymmetric Noise and 33 Noise-Free trials), corresponding to 12 excluded trials. For interpersonal hand movement coordination, 104 trials were retained (70 Noise and 34 Noise-Free), corresponding to 13 excluded trials. For intrapersonal hand movement coordination, 110 trials were retained (74 Noise and 36 Noise-Free), corresponding to 7 excluded trials.

Consequently, sample sizes differ slightly across analyses according to the availability of the signals required for each measure. A summary of data retention is provided in Table S1.

**Table S1. Data retention across analyses. Number of conversation trials retained and excluded from each analysis following the general dyad-level exclusion and analysis-specific data-quality assessment.**

| Analysis | Available trials | Retained | Excluded | Condition breakdown |
|---|---|---|---|---|
| Subjective ratings | 117 | 117 | 0 | 78 Noise / 39 Noise-Free |
| Turn-taking | 117 | 117 | 0 | 78 Noise / 39 Noise-Free |
| Interpersonal head coordination | 117 | 105 | 12 | 72 Noise / 33 Noise-Free |
| Interpersonal hand coordination | 117 | 104 | 13 | 70 Noise / 34 Noise-Free |
| Intrapersonal hand coordination | 117 | 110 | 7 | 74 Noise / 36 Noise-Free |

For intrapersonal analyses, each retained conversation could contribute one observation per participant when sufficient bilateral hand-movement data were available. Therefore, the number of participant-level observations used in the intrapersonal analysis differs from the number of retained conversation trials reported above.

**S2. Turn-taking annotation and feature extraction.** **Conversational unit definitions**

Turn-taking structure was annotated using two primary interactional units. A *turn* was defined as a continuous stretch of speech produced by one participant before yielding the conversational floor. A *backchannel* was defined as a brief listener-produced utterance (e.g., “uh-huh”, “yeah”, “okay”) that was relevant to the ongoing turn but did not constitute an attempt to take the conversational floor. These definitions were applied throughout automated classification and subsequent manual correction.

#### Automated annotation pipeline

Speech annotations were generated using an automated pipeline (1) combining voice activity detection, cross-talk filtering, automatic transcription, and classification of conversational units. The pipeline is publicly available at: https://github.com/haraldsr/Speech_VAD_Diarization_Transcription.

Speech activity was first detected separately for each participant’s audio channel using the rVAD algorithm at a temporal resolution of 10 ms. To reduce cross-talk between the two simultaneously recorded channels, detected segments were subsequently filtered using speaker-specific root-mean-square (RMS) energy.

Temporally adjacent speech segments from the same speaker were merged when separated by pauses shorter than 500 ms, thereby reducing fragmentation caused by short within-turn pauses. The resulting candidate units were transcribed using the Whisper-large-v3 automatic speech recognition model.

Candidate units were classified as turns or backchannels using Shannon entropy of their transcribed content, with lower lexical variability used to identify candidate backchannels. A subsequent rule-based refinement step incorporated the temporal context of each segment. Consecutive turns from the same speaker were merged, candidate backchannels occurring outside another participant’s turn were reclassified as turns, and short turns embedded within the other participant’s ongoing turn were reclassified as backchannels.

#### Manual correction

All automatically generated annotations were subsequently manually reviewed. A total of 117 conversations were inspected, with corrections made to speech onsets and offsets and to turn/backchannel classifications where necessary.

Manual correction was performed using ELAN (Figure S1), allowing annotations to be evaluated against the corresponding audio, transcription, timing.

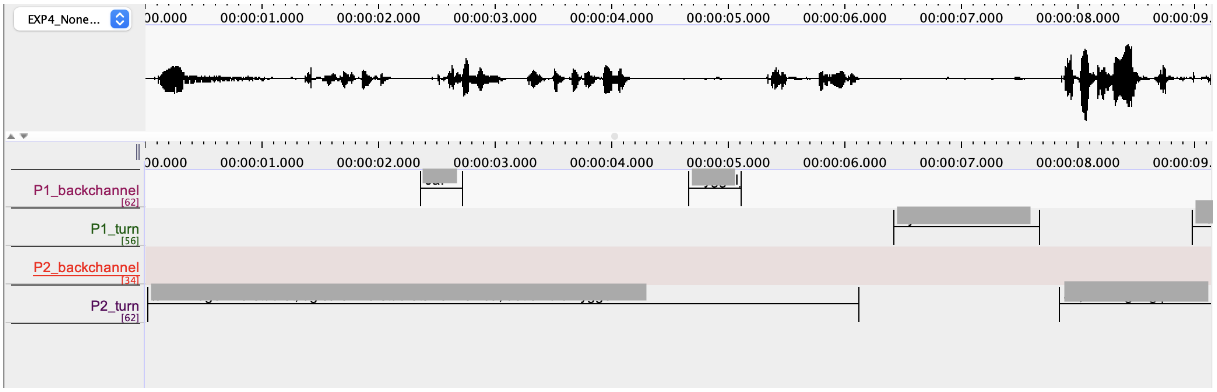


**Fig. S1. Example of manual turn-taking annotation in ELAN.** Automatically generated speech segments were manually reviewed and corrected with respect to segment boundaries and turn/backchannel classification. Transcriptions are obscured here due to privacy.

### Extraction of turn-taking measures

Four measures used in the main analyses were derived from the corrected annotations: average turn duration, speaker contribution, backchannel frequency, and floor-transfer offset (FTO).

Average turn duration was calculated separately for each participant as the mean duration of their annotated turns within each conversation. Speaker contribution was calculated as the proportion of the dyad's combined speaking time contributed by each participant. Backchannel frequency was defined as the total number of annotated backchannels produced by each participant during a conversation.

FTOs were calculated between consecutive turns produced by different speakers as the temporal difference between the offset of the preceding turn and the onset of the following turn. Positive FTOs therefore represent silent gaps between turns, whereas negative FTOs represent temporal overlap between speakers. Backchannels were excluded because they did not constitute attempts to take the conversational floor. Turns fully embedded within another speaker's turn were likewise excluded because they did not represent a floor-transfer event.

For participant-level analyses, FTOs were assigned to the participant initiating the subsequent turn. For the analysis relating turn-transition timing to conversational enjoyment, valid FTOs from both participants were pooled within each conversation and averaged to obtain a single trial-level measure.

### Example conversational structure

Figure S2 illustrates the resulting annotations from a representative 10-minute conversation.

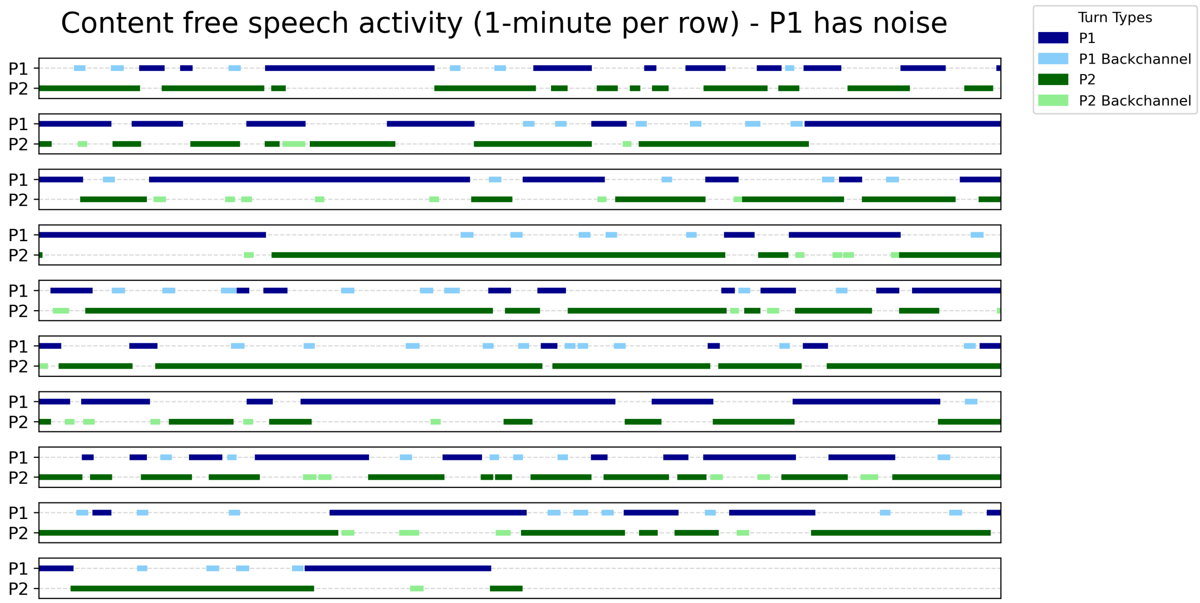


**Fig. S2. Example annotated turn-taking structure from a representative 10-minute conversation.** Each row represents one minute of interaction. Segments show speech activity from the two participants and the classification of conversational units as turns or backchannels.

### S3. Personality traits. Trait Extraversion and Post-Conversation Subjective Experience

To examine whether individual differences in personality moderated subjective conversational outcomes under asymmetric noise, we tested the effect of trait Extraversion across all three post-conversation measures: conversational flow ($z$-scored), conversational enjoyment ($z$-scored), and interpersonal closeness (Inclusion of Other in the Self [IOS] scale).

Conversational flow and enjoyment were modeled using linear mixed-effects models with listening condition (*Noise-Self*, *Noise-Other*, *Noise-Free*), mean-centered Extraversion, and their interaction entered as fixed effects, and nested random intercepts for participants within dyads (1 | Dyad/Participant). Interpersonal closeness (IOS) was modeled using a cumulative link mixed model (CLMM) for ordinal ratings with a random intercept for dyad.

As shown in Supplementary Figure S3, individual Extraversion did not significantly predict perceived conversational flow ($F(1, 5.00) = 0.14$, $p = .722$), conversational enjoyment ($F(1, 5.00) = 0.02$, $p = .898$), or interpersonal closeness ($\chi^2(1) = 0.14$, $p = .710$). Furthermore, the interaction between listening condition and Extraversion was non-significant across all three measures (flow: $F(2, 31.00) = 0.99$, $p = .384$; enjoyment: $F(2, 31.00) = 1.91$, $p = .165$; IOS: $\chi^2(2) = 0.63$, $p = .731$). These results indicate that subjective conversational quality and interpersonal closeness were unaffected by extraversion across acoustic conditions.

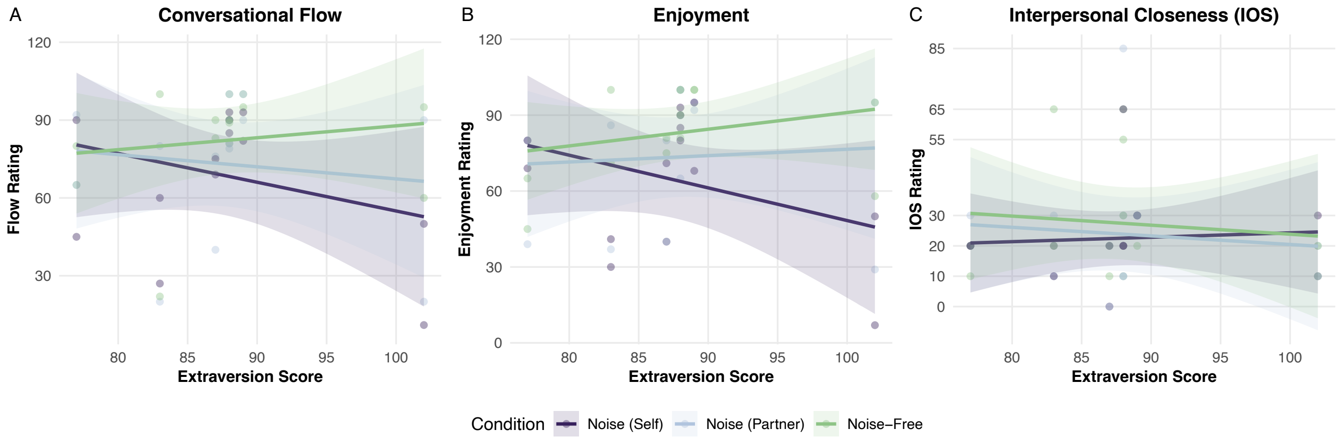


**Fig. S3. Relationship between trait Extraversion and subjective conversational ratings across listening conditions.** (A) Perceived conversational flow, (B) conversational enjoyment, and (C) interpersonal closeness (IOS) plotted as a function of individual Extraversion scores across Noise-Self, Noise-Other, and Noise-Free conditions. Solid lines represent linear regression fits with 95% confidence intervals. Across all measures, Extraversion did not significantly predict subjective outcomes or interact with acoustic condition.

#### Trait Extraversion and Verbal Turn-Taking Features

To evaluate whether individual differences in personality moderated conversational coordination under asymmetric acoustic noise, we examined the relationship between trait Extraversion and four core turn-taking parameters: average turn duration, speaker contribution (percentage of total speaking time), backchannel frequency, and floor-transfer offset (FTO).

Separate linear mixed-effects models were fitted for each measure with listening condition (*Noise-Self*, *Noise-Other*, *Noise-Free*), mean-centered Extraversion, and their interaction specified as fixed effects, alongside a random intercept for dyad (1 | EXP_ID). Fixed effects were evaluated using Type III ANOVA with Satterthwaite-approximated degrees of freedom.

As shown in Supplementary Figure S4, trait Extraversion showed a significant main effect on speaker contribution ($F(1, 219) = 5.84$, $p = .017$), indicating that more extraverted individuals contributed a larger proportion of the dyad's total speaking time across listening conditions. Extraversion did not significantly predict average turn duration ($F(1, 217.90) = 1.08$, $p = .299$), backchannel count ($F(1, 214.13) = 1.15$, $p = .286$), or floor-transfer offset ($F(1, 209.90) = 2.91$, $p = .090$).

Crucially, the interaction between Extraversion and listening condition was non-significant across all four turn-taking features (turn duration: $F(2, 182.37) = 0.50$, $p = .608$; speaker contribution: $F(2, 219) = 1.50$, $p = .225$; backchannel count: $F(2, 181.98) = 1.55$, $p = .216$; FTO: $F(2, 182.33) = 0.36$, $p = .696$). Thus, although more extraverted individuals contributed more of the dyad's overall speaking time, there was no evidence that trait Extraversion moderated the effects of asymmetric noise on verbal turn-taking dynamics.

### S4. Movement preprocessing and recurrence analysis. Pre-processing details

Motion capture data were acquired at 100 Hz using a Vicon tracking system. Three-dimensional center-of-mass (COM) trajectories were reconstructed for each effector by averaging the corresponding marker positions: six markers for the head and two markers for each hand.

Missing data were assessed prior to interpolation. Across recordings, the proportion of missing samples was low (mean = 0.36%, maximum = 4.29% per participant). Missing samples were linearly interpolated to preserve temporal continuity. Movement signals were converted from millimetres to metres and temporally aligned. Signals were low-pass filtered using a 4th-order Butterworth filter with a 2 Hz cutoff and subsequently downsampled from 100 Hz to 10 Hz using the Largest-Triangle-Three-Buckets (LTTB) algorithm.

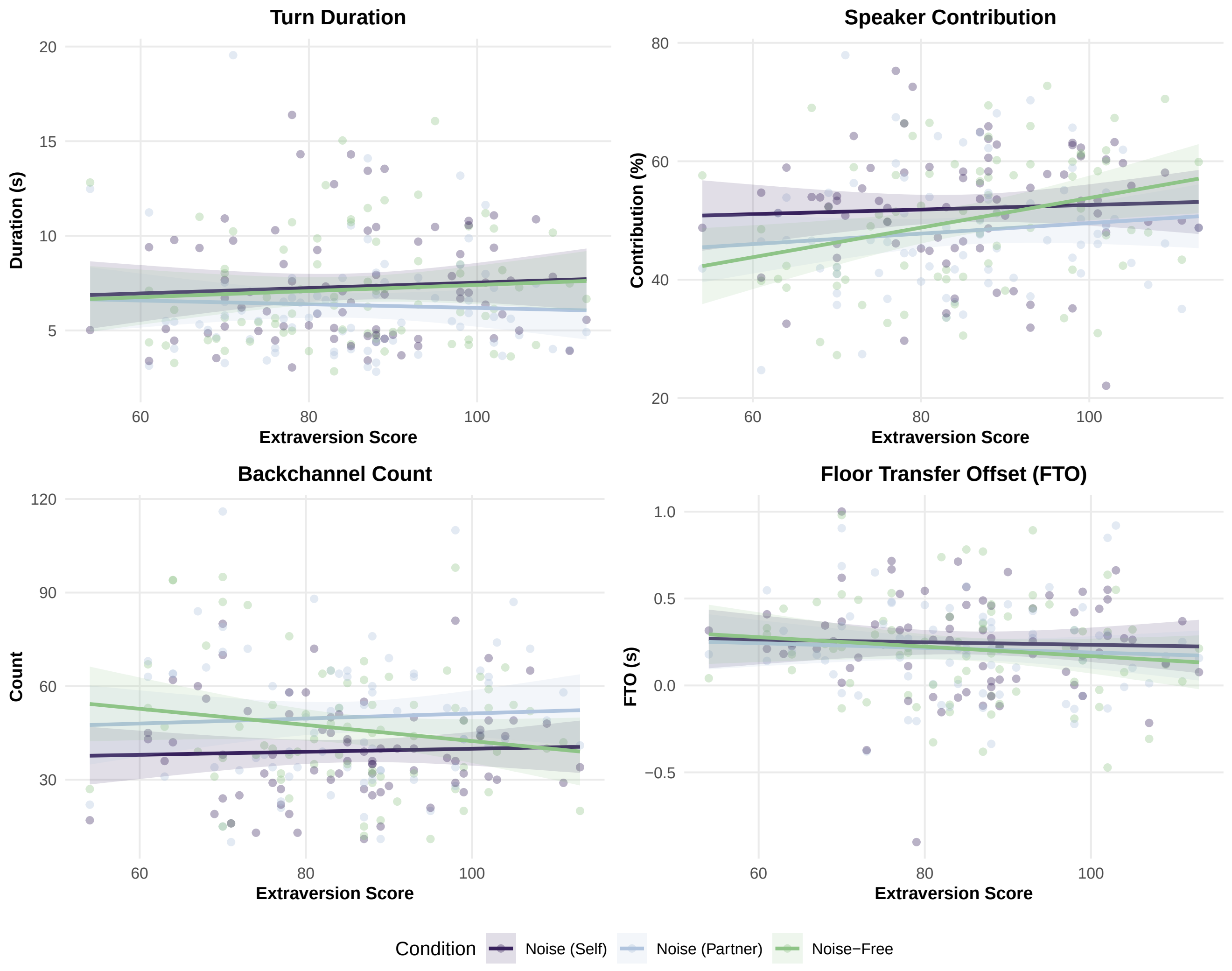


**Fig. S4. Relationship between trait Extraversion and verbal turn-taking features across listening conditions.** (A) Average turn duration, (B) speaker contribution, (C) backchannel count, and (D) floor-transfer offset (FTO) plotted as a function of individual Extraversion scores for Noise-Self, Noise-Other, and Noise-Free conditions. Solid lines represent linear model fits with 95% confidence intervals.

Analyses were conducted on a standardized 10-minute segment from each trial, beginning after an initial 30-s period. Example preprocessed movement time series are shown for a representative dyad in Figures S5 and S6.

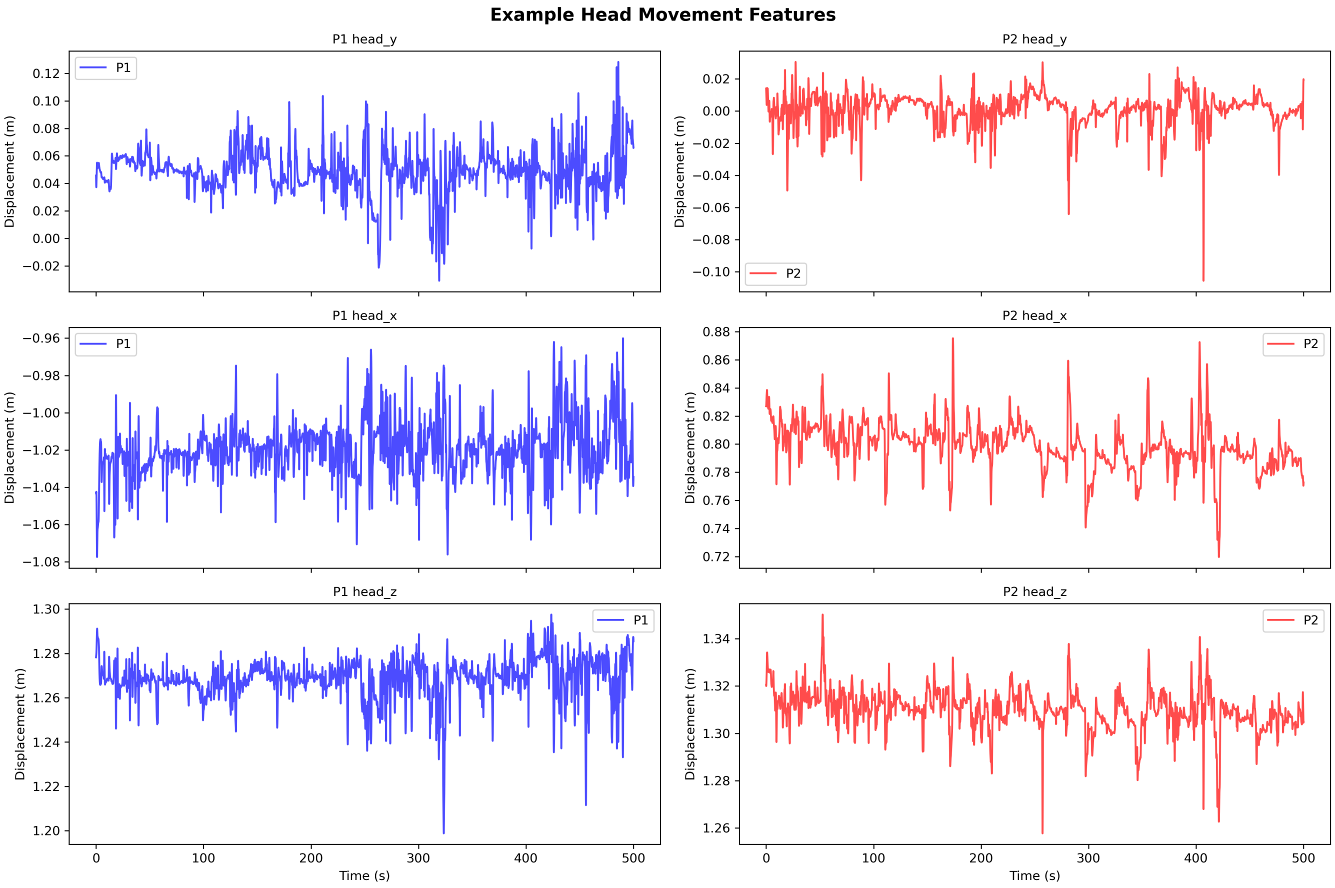


**Fig. S5. Example preprocessed head movement signals.** Head center-of-mass (COM) trajectories for a representative dyad over the analysis window following interpolation, filtering, and downsampling.

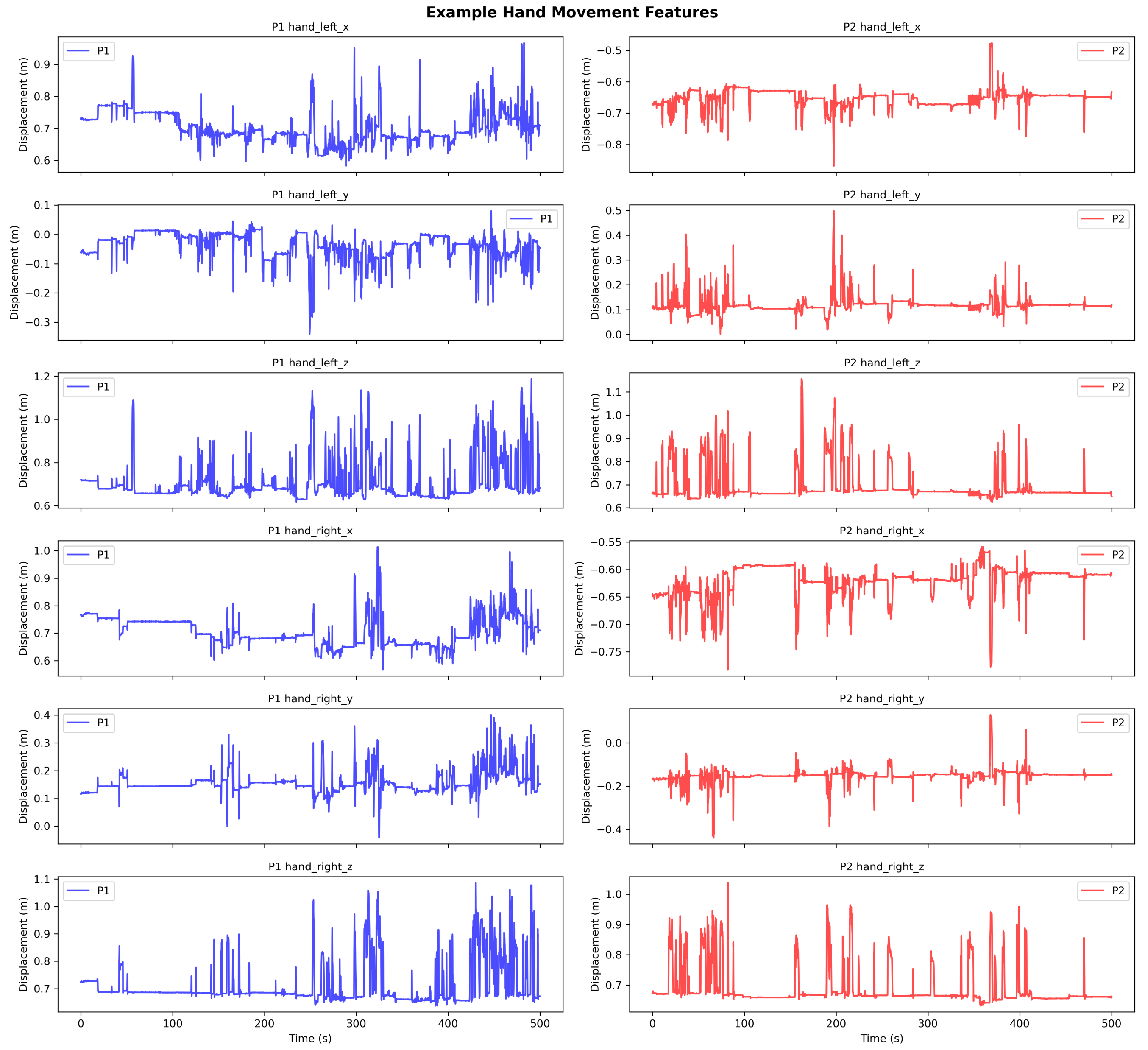


**Fig. S6. Example preprocessed hand movement signals.** Hand center-of-mass (COM) trajectories for a representative dyad over the analysis window following interpolation, filtering, and downsampling.

### Construction of multivariate movement signals

For interpersonal head coordination, each participant was represented by a three-dimensional time series containing the $x$, $y$, and $z$ trajectories of the head COM. For interpersonal hand coordination, the trajectories of the left and right hands were combined to represent each participant's hand movement. Because participants were seated facing one another, P2 movement signals were mirrored along the $y$-axis before interpersonal analysis to establish a consistent spatial reference frame across partners.

For asymmetric noise trials, participant signals were subsequently reordered such that the noise-exposed participant was always assigned as the first (reference) time series and the noise-free partner as the second time series. This ensured a consistent interpretation of directional temporal lags across dyads and trials. For intrapersonal hand coordination, left- and right-hand movement trajectories were compared within each participant using their corresponding $x$, $y$, and $z$ coordinates. This provided a measure of within-person coordination between the two hands. All movement dimensions were z-standardized before recurrence analysis.

### MDCRQA parameter estimation

Interpersonal and intrapersonal movement coordination were quantified using multidimensional cross-recurrence quantification analysis (MDCRQA)(2). MDCRQA identifies recurrent states between multivariate movement signals based on their proximity within the reconstructed state space.

Embedding parameters were estimated separately for each trial using a data-driven optimization procedure. Embedding dimension ($m$), delay ($\tau$), and recurrence radius ($r$) were selected while targeting a recurrence rate within the 2.9-3.1% range. The radius was searched within a constrained range to maintain comparable recurrence density across trials and conditions.

A summary of the resulting parameters is provided in Table S2.

**Table S2. MDCRQA parameter optimization across movement analyses. Optimized embedding dimension ($m$), radius ($r$), and delay ($\tau$) are reported as mean $\pm$ SD across trials.**

| Level | Condition | n | Mean $m$ | SD $m$ | Mean $r$ | SD $r$ | Mean $\tau$ | SD $\tau$ |
|---|---|---|---|---|---|---|---|---|
| Interpersonal | Head (Noise) | 72 | 2.38 | 1.23 | 0.32 | 0.04 | 3.86 | 1.03 |
| Interpersonal | Head (None) | 33 | 2.24 | 0.87 | 0.32 | 0.03 | 3.61 | 0.66 |
| Interpersonal | Hand (Noise) | 70 | 2.57 | 1.19 | 0.36 | 0.07 | 6.06 | 2.73 |
| Interpersonal | Hand (None) | 34 | 2.79 | 1.45 | 0.38 | 0.05 | 6.00 | 2.75 |
| Intrapersonal | Hand (Noise_Other) | 68 | 2.88 | 1.24 | 0.24 | 0.07 | 5.54 | 3.47 |
| Intrapersonal | Hand (Noise_Self) | 68 | 3.15 | 1.18 | 0.24 | 0.06 | 5.85 | 3.06 |
| Intrapersonal | Hand (None) | 69 | 3.06 | 1.21 | 0.26 | 0.06 | 5.26 | 3.17 |

The minimum diagonal and vertical line lengths were fixed at 20 samples throughout the analyses. All other analysis settings were held constant across experimental conditions.

Figure S7 shows an example cross-recurrence plot for interpersonal head movement coordination.

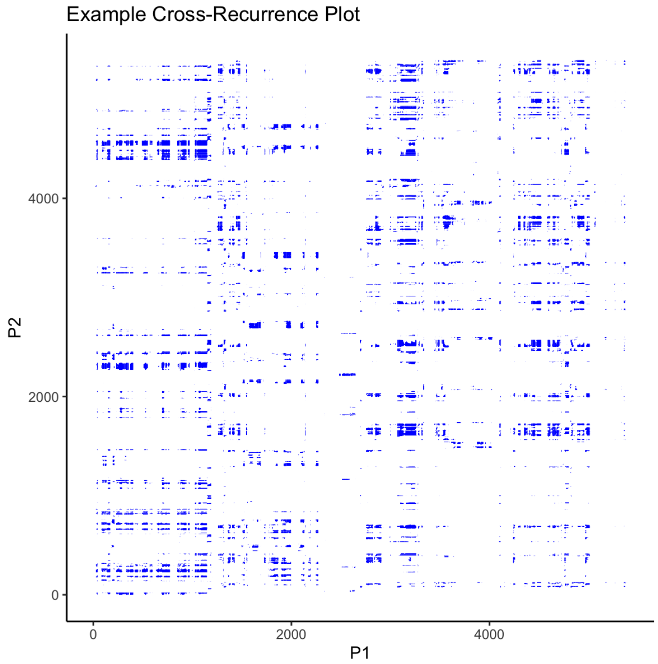


**Fig. S7. Example cross-recurrence plot for interpersonal head movement coordination.** Recurrence points indicate recurrent movement states between the two participants over time.

### Diagonal cross-recurrence profiles

The temporal structure of recurrence was quantified using diagonal cross-recurrence profiles (DCRP) (2). DCRP summarizes the proportion of recurrent states along diagonals of the cross-recurrence plot, providing recurrence as a function of temporal lag.

For interpersonal head and hand coordination, DCRPs were calculated over a lag window of $-5$ to $+5$ s. Because asymmetric trials were oriented with the noise-exposed participant as the reference time series, negative lags indicate temporal precedence of movements from the noise-exposed participant, whereas positive lags indicate temporal precedence of movements from the noise-free partner.

In the Noise-Free condition, P1/P2 assignment was arbitrary and therefore provided no meaningful directional reference. To prevent arbitrary participant assignment from introducing directional structure into the condition-level profile, each Noise-Free DCRP was symmetrized by averaging the observed profile with its time-reversed counterpart:

$$DCRP_{\text{sym}}(l) = \frac{DCRP(l) + DCRP(-l)}{2}. \quad (145)$$

 

This procedure preserves the magnitude and lag-dependent structure of recurrence while accounting for arbitrary participant labeling.

For intrapersonal hand coordination, DCRPs were calculated between the left- and right-hand movement signals over the same $-5$ to $+5$ s lag window. For statistical analysis, recurrence values were averaged within 1-s signed lag bins, retaining temporal information without assuming symmetry around zero lag.

#### Surrogate DCRP analysis

To estimate recurrence expected in the absence of temporal coupling, surrogate DCRP profiles were generated by randomly permuting the temporal order of one participant's movement signal (3). For each trial, the temporal samples of each dimension of the second participant's multivariate time series were independently randomly permuted. This preserved the distribution of values within each movement dimension while disrupting its temporal organization and temporal relationship with the first participant.

The shuffled signal was then entered into the same DCRP procedure as the empirical data using the corresponding analysis parameters. This procedure was repeated 19 times for each trial, and the resulting DCRP profiles were averaged at each temporal lag to obtain a trial-level surrogate profile. The surrogate therefore represents recurrence expected when the observed movement states are retained but their temporal organization is removed.

Figure S8 shows an empirical DCRP and its corresponding surrogate baseline for a representative asymmetric-noise trial.

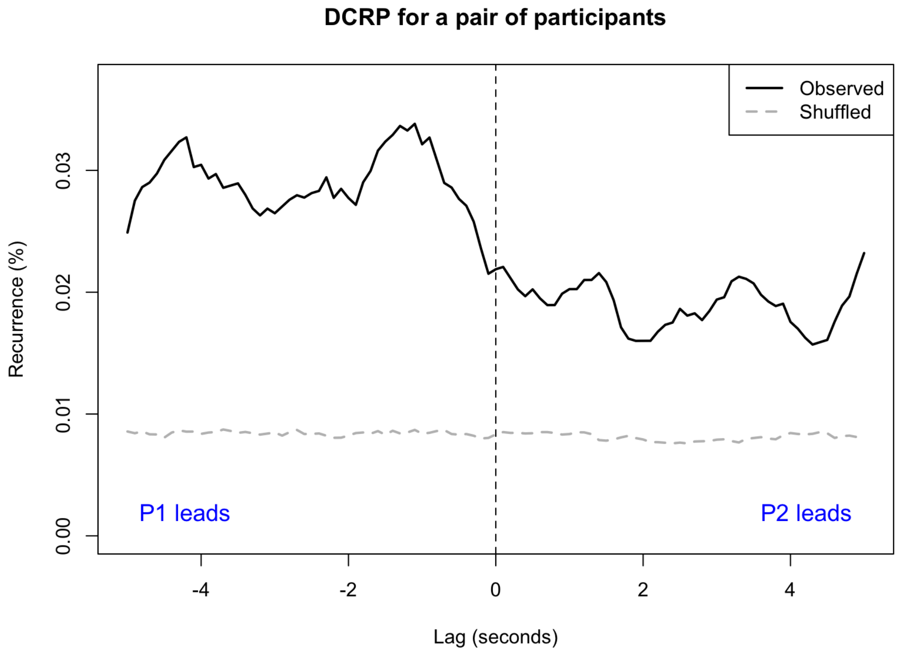


**Fig. S8. Example diagonal cross-recurrence profile (DCRP) for interpersonal head movement coordination.** The noise-exposed participant is assigned as the reference participant (P1). Negative lags indicate P1's movement preceded P2, whereas positive lags indicate P2's movement preceded P1. The black line shows the empirical DCRP and the dashed line the temporally randomized surrogate baseline. The vertical line indicates zero lag.

Surrogate profiles are presented as a chance-level reference and were omitted from the main figures to emphasize differences in the temporal structure of observed coordination across experimental conditions.

**S5. Control analyses of movement magnitude.** To assess whether the observed movement coordination effects could be explained by differences in overall movement magnitude across listening conditions, we conducted additional control analyses based on mean movement speed. Movement magnitude was calculated from three-dimensional displacement between consecutive samples and converted to speed based on the 10-Hz sampling rate. We first tested whether movement magnitude differed across listening conditions and then repeated each recurrence analysis with movement magnitude included as a covariate.

#### Interpersonal head movement

For interpersonal head coordination, movement magnitude was calculated at the conversation level as the mean three-dimensional head movement speed across both participants. Mean head movement speed did not significantly differ between Noise and Noise-Free conversations ($F(1, 66.43) = 2.01$, $p = .161$; Figure S9A).

We then repeated the interpersonal head recurrence analysis with mean head movement speed included as a covariate. Mean head speed was not significantly associated with recurrence ($F(1, 72.90) = 1.02$, $p = .317$). The Condition $\times$ lag interaction remained significant after accounting for movement magnitude ($F(1, 10498) = 55.14$, $p < .001$), while the main effect of condition remained non-significant ($F(1, 70.20) = 0.92$, $p = .340$). The Condition × lag coefficient was unchanged between the original and movement-adjusted models ($\beta = -2.33 \times 10^{-4}$ in both models).

Thus, accounting for mean head movement speed did not alter the condition-dependent temporal pattern of interpersonal head coordination.

**Interpersonal hand movement**

For interpersonal hand coordination, movement magnitude was calculated at the conversation level as the mean three-dimensional movement speed across the left and right hands of both participants. Mean hand movement speed was significantly greater in Noise than in Noise-Free conversations ($F(1, 66.04) = 4.66$, $p = .035$; Figure S9B), with an estimated difference of 0.0077 m/s ($SE = 0.0036$).

We next repeated the interpersonal hand recurrence analysis with mean hand movement speed included as a covariate. Mean hand speed was not significantly associated with recurrence ($F(1, 101) = 0.57$, $p = .451$). The Condition × lag interaction remained non-significant ($F(1, 10398) = 2.74$, $p = .098$), with an identical interaction coefficient before and after adjustment ($\beta = 4.92 \times 10^{-5}$). The main effect of condition also remained non-significant ($F(1, 101) = 0.75$, $p = .388$).

Thus, although hand movement magnitude was greater during Noise conversations, accounting for this difference did not alter the interpersonal hand recurrence results.

**Intrapersonal hand movement**

For intrapersonal hand coordination, movement magnitude was calculated separately for each participant as the mean three-dimensional movement speed across their left and right hands. Mean hand movement speed differed across listening conditions ($F(2, 95.53) = 3.17$, $p = .046$; Figure S9C). Estimated marginal means were 0.0741 m/s ($SE = 0.0046$) in *Noise-Free*, 0.0813 m/s ($SE = 0.0046$) in *Noise-Self*, and 0.0839 m/s ($SE = 0.0045$) in *Noise-Other*. Tukey-adjusted comparisons showed significantly greater movement magnitude in *Noise-Other* than in *Noise-Free* ($\beta = 0.0097$ m/s, $p = .044$), while the other pairwise comparisons were not significant (all $p \geq .173$).

We then repeated the intrapersonal recurrence analysis with mean hand movement speed included as a covariate. Mean hand speed was not significantly associated with recurrence ($F(1, 145.13) = 0.13$, $p = .722$), while the effect of listening condition remained significant ($F(2, 136.21) = 3.69$, $p = .027$). The condition contrasts were also essentially unchanged after adjustment. Recurrence remained higher in *Noise-Self* than in *Noise-Other* ($\beta = 0.0144$, $p = .049$), while the *Noise-Self* versus *Noise-Free* ($\beta = 0.0137$, $p = .061$) and *Noise-Other* versus *Noise-Free* ($\beta = -0.0006$, $p = .994$) contrasts remained non-significant.

Thus, although hand movement magnitude varied across listening conditions, accounting for mean hand movement speed did not alter the condition effect on intrapersonal hand recurrence.

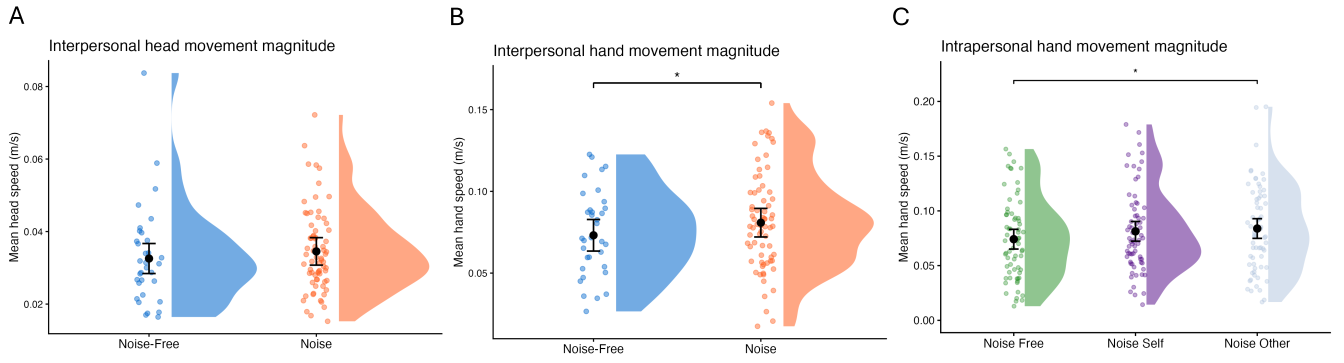


**Fig. S9. Movement magnitude across listening conditions.** Mean three-dimensional movement speed for (A) interpersonal head movement, calculated at the conversation level across both participants; (B) interpersonal hand movement, calculated at the conversation level across both hands and participants; and (C) intrapersonal hand movement, calculated separately for each participant across the left and right hands. Points show individual observations and distributions show the observed data; superimposed estimates indicate estimated marginal means with 95% confidence intervals.

**A. S6. Backchannel rate analysis.** As an additional analysis of backchanneling, we directly calculated each participant's backchannel rate relative to the amount of partner speech available. Backchannel rate was defined as the number of backchannels produced per minute of interaction-partner speaking time:

$$R_{ij} = \frac{B_{ij}}{T_{ij}^{\text{partner}}/60}, \quad [1]$$

where $B_{ij}$ represents the number of backchannels produced by participant $i$ and $T_{ij}^{\text{partner}}$ represents the total speaking duration of their interaction partner in seconds. This provides a direct measure of backchannel production while accounting for differences in the opportunity to backchannel.

Backchannel rate was analyzed using a linear mixed-effects model with individual listening condition (*Noise-Self*, *Noise-Other*, *Noise-Free*) as a categorical fixed effect and random intercepts for participant and dyad. The effect of condition was evaluated using Type III ANOVA with Satterthwaite-approximated degrees of freedom, followed by Tukey-adjusted estimated marginal mean (EMM) comparisons.

Backchannel rate differed significantly across listening conditions, $F(2, 154) = 12.13$, $p < .001$ (Figure S10). Noise-exposed participants produced fewer backchannels per minute of partner speech ($M = 9.25$, $SE = 0.42$) than their non-exposed partners ($M = 10.77$, $SE = 0.42$; $b = -1.52$, $SE = 0.32$, $t = -4.79$, $p < .001$). Noise-exposed participants also showed a lower backchannel rate than participants in the *Noise-Free* condition ($M = 10.32$, $SE = 0.42$; $b = -1.08$, $SE = 0.32$, $t = -3.39$, $p = .003$). In contrast, backchannel rates did not differ significantly between non-exposed participants and participants in the *Noise-Free* condition ($b = 0.45$, $SE = 0.32$, $t = 1.40$, $p = .342$). Thus, directly normalizing backchannel counts by partner speaking time showed the same pattern of condition differences.

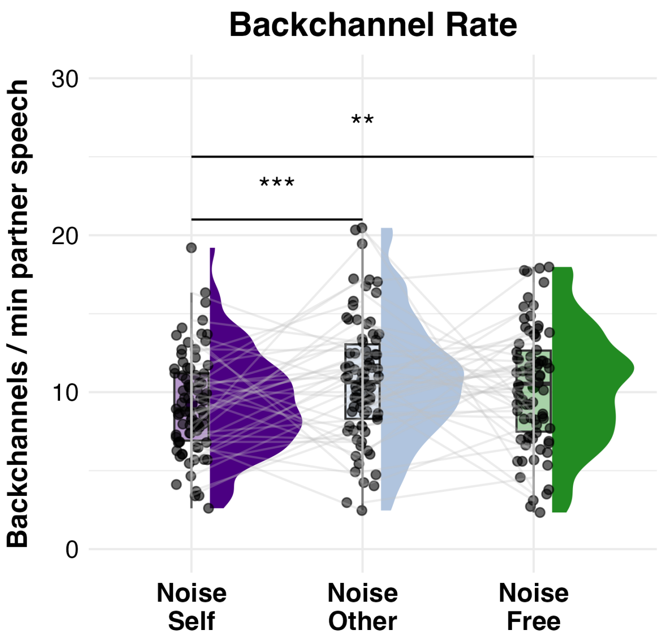


**Fig. S10. Backchannel rate across listening conditions.** Backchannel rate was calculated as the number of backchannels produced per minute of interaction-partner speaking time. Points show participant-level observations across the *Noise-Self*, *Noise-Other*, and *Noise-Free* conditions. Significance annotations indicate Tukey-adjusted pairwise comparisons.

**S7. Statistical analysis.** All statistical analyses were conducted using mixed-effects models to account for the repeated-measures structure of the data. Linear mixed-effects models were estimated using restricted maximum likelihood (REML), with significance of fixed effects assessed using Type III analysis of variance with Satterthwaite-approximated degrees of freedom unless otherwise specified. Post hoc comparisons were based on estimated marginal means (EMMs). Detailed test statistics for the primary analyses are reported in the main Results; the sections below provide the complete model specifications and additional analyses required for interpretation and replication.

### Subjective ratings

Conversational flow and enjoyment were analyzed separately using linear mixed-effects models. In both analyses, individual participant ratings were the unit of observation and listening condition was entered as a categorical fixed effect with three levels: *Noise-Self*, in which the participant was exposed to noise; *Noise-Other*, in which their partner was exposed to noise; and *Noise-Free*, in which neither participant was exposed to noise. Random intercepts were included for dyad and trial nested within dyad. The models were specified as:

$$\text{Rating}_{ijk} = \beta_0 + \beta_1 \text{Condition}_{ijk} + u_{\text{Dyad}_j} + u_{\text{Trial}_{k(j)}} + \epsilon_{ijk}, \quad [2]$$

where Rating represents either conversational flow or enjoyment. Significant condition effects were followed by pairwise comparisons of EMMs with Tukey adjustment for multiple comparisons. Cohen's $d$ was calculated from the model-estimated contrasts using the residual standard deviation of the corresponding model.

Interpersonal closeness was measured using the seven-level Inclusion of Other in the Self (IOS) scale and was therefore analyzed using a cumulative link mixed model (CLMM). The model included trial number as a continuous predictor, listening condition, and their interaction, with a random intercept for dyad:

$$\text{logit}\left[P(\text{IOS}_{ij} \le c)\right] = \theta_c - \left[\beta_1 \text{Trial}_{ij} + \beta_2 \text{Condition}_{ij} + \beta_3 (\text{Trial} \times \text{Condition})_{ij} + u_{\text{Dyad}_j}\right], \quad [3]$$

where $\theta_c$ represents the threshold associated with each ordinal cutpoint. Significance of fixed effects was assessed using Type III Wald omnibus tests ($\chi^2$).

#### Turn-taking measures

Effects of asymmetric listening condition on turn-taking were assessed separately for average turn duration, speaker contribution, backchanneling, and floor-transfer offset (FTO). Participant-level observations were categorized as *Noise-Self*, *Noise-Other*, or *Noise-Free*. Speaker contribution was calculated as the percentage of total dyadic speaking time attributable to each participant, based on the summed duration of their speaking turns.

Average turn duration, speaker contribution, and FTO were analyzed using linear mixed-effects models with listening condition as a categorical fixed effect and random intercepts for participant and dyad:

$$Y_{ij} = \beta_0 + \beta_1 \text{Condition}_{ij} + u_{\text{Dyad}_j} + u_{\text{Participant}_i} + \epsilon_{ij}, \quad [4]$$

where $Y$ represents the respective turn-taking measure. Fixed effects were evaluated using Type III ANOVA with Satterthwaite-approximated degrees of freedom. Significant condition effects were followed by Tukey-adjusted pairwise EMM comparisons. Cohen's $d$ was calculated from the estimated contrasts using the residual standard deviation of the respective model.

Because the number of backchannels depends on the amount of partner speech available as an opportunity to backchannel, backchannel counts were analyzed separately using a negative-binomial mixed-effects model. Listening condition was entered as a categorical fixed effect, with random intercepts for participant and dyad, and the interaction partner's total speaking duration was included as an exposure offset on the log scale:

$$\log\left(E[B_{ij}]\right) = \beta_0 + \beta_1 \text{Condition}_{ij} + \log(T_{ij}^{\text{partner}}) + u_{\text{Dyad}_j} + u_{\text{Participant}_i}, \quad [5]$$

where $B_{ij}$ represents the number of backchannels produced by participant $i$ and $T_{ij}^{\text{partner}}$ represents the total speaking duration of their interaction partner. The model therefore estimates condition differences in backchanneling relative to the amount of partner speech available. The overall effect of condition was evaluated using a Type III Wald $\chi^2$ test, followed by Tukey-adjusted pairwise comparisons of estimated rate ratios.

#### Relationship between floor-transfer offset and enjoyment

To examine whether turn-transition timing was related to subjective conversational experience, valid FTOs from both participants were pooled within each conversation and averaged to obtain a trial-level measure of turn-transition timing. Enjoyment ratings from the two participants were similarly averaged to obtain a dyad-level enjoyment score for each conversation.

Mean enjoyment was modeled as a function of mean FTO, listening condition (*Noise* vs. *Noise-Free*), and their interaction, with a random intercept for dyad:

$$\text{Enjoyment}_i = \beta_0 + \beta_1 \text{FTO}_i + \beta_2 \text{Condition}_i + \beta_3 (\text{FTO}_i \times \text{Condition}_i) + u_{\text{Dyad}} + \epsilon_i. \quad [6]$$

The FTO × Condition interaction tested whether the relationship between turn-transition timing and enjoyment differed between asymmetric-noise and noise-free conversations.

#### Interpersonal movement coordination

Interpersonal head and hand movement coordination were analyzed separately using the lag-resolved DCRP recurrence values. For each movement modality, recurrence was modeled as a function of listening condition (*Noise* vs. *Noise-Free*), temporal lag, and their interaction:

$$\text{Recurrence}_i = \beta_0 + \beta_1 \text{Condition}_i + \beta_2 \text{Lag}_i + \beta_3 (\text{Condition}_i \times \text{Lag}_i) + u_{\text{Dyad}} + u_{\text{Trial}} + \epsilon_i. \quad [7]$$

Temporal lag was entered as a continuous predictor ranging from $-5$ to $+5$ s. Random intercepts for dyad and trial accounted for the repeated recurrence observations obtained across temporal lags. The Condition × Lag interaction was of primary interest because it tested whether the temporal structure of interpersonal coordination differed between asymmetric-noise and noise-free conversations.

For head movements, the significant difference between positive and negative lag recurrences was further examined by testing the symmetry of the DCRP around zero lag. Continuous lag values were grouped into 1-s bins, and each negative-lag bin was contrasted with its corresponding positive-lag bin within each listening condition. For example, recurrence between $-3$ and $-2$ s was compared with recurrence between $+2$ and $+3$ s. Because the noise-exposed participant was consistently assigned as the reference participant, greater recurrence at negative than corresponding positive lags indicates greater temporal precedence of the noise-exposed participant's movements.

Contrasts were calculated from EMMs using asymptotic $z$-tests, with False Discovery Rate (FDR) correction applied separately across the five lag symmetry contrasts within each condition. Complete results are reported in Table S3.

**Table S3. Symmetry contrasts for interpersonal head movement DCRP. Estimated differences in recurrence between corresponding negative and positive temporal lag bins within the Noise-Free and asymmetric Noise conditions. Positive estimates indicate greater recurrence at the negative lag than at the corresponding positive lag. $p$-values are FDR-adjusted within condition.**

| Condition | Contrast | Estimate | SE | $z$ | $p_{\mathrm{FDR}}$ |
|---|---|---|---|---|---|
| Noise-Free | $[-5,-4]-[4,5]$ | 0.000016 | 0.000332 | 0.048 | .9620 |
| | $[-4,-3]-[3,4]$ | 0.000100 | 0.000340 | 0.294 | .9620 |
| | $[-3,-2]-[2,3]$ | -0.000103 | 0.000340 | -0.303 | .9620 |
| | $[-2,-1]-[1,2]$ | -0.000021 | 0.000340 | -0.061 | .9620 |
| | $[-1,0]-[0,1]$ | -0.000043 | 0.000340 | -0.126 | .9620 |
| Noise | $[-5,-4]-[4,5]$ | 0.00186 | 0.000225 | 8.287 | $<.0001$ |
| | $[-4,-3]-[3,4]$ | 0.00154 | 0.000230 | 6.697 | $<.0001$ |
| | $[-3,-2]-[2,3]$ | 0.00157 | 0.000230 | 6.836 | $<.0001$ |
| | $[-2,-1]-[1,2]$ | 0.000871 | 0.000230 | 3.789 | .0002 |
| | $[-1,0]-[0,1]$ | 0.000855 | 0.000230 | 3.719 | .0002 |

The Noise-Free profile showed no evidence of directional asymmetry, with all five corresponding negative–positive lag contrasts close to zero and non-significant after FDR correction. In contrast, all five contrasts were positive and significant in the asymmetric Noise condition, indicating systematically greater recurrence at negative than at corresponding positive lags across the examined temporal window.

### Intrapersonal hand movement coordination

Intrapersonal hand coordination was quantified from the lag-resolved DCRP between each participant's left- and right-hand movements. To retain temporal information without assuming symmetry around zero lag, recurrence values were averaged within 1-s signed lag bins from $-5$ to $+5$ s.

Binned recurrence was analyzed using a linear mixed-effects model with listening condition (*Noise-Self*, *Noise-Other*, *Noise-Free*) and lag bin as fixed effects. Random intercepts were included for participant, conversation, and participant-specific DCRP profile. These accounted respectively for repeated observations from the same participant across conversations, the shared interaction between the two partners within a conversation, and repeated lag bins within each participant-specific profile:

$$\mathrm{Recurrence}_{ijkl} = \beta_0 + \beta_1 \mathrm{Condition}_i + \gamma_j \mathrm{LagBin}_j + u_{\mathrm{Participant}} + u_{\mathrm{Conversation}} + u_{\mathrm{Profile}} + \epsilon_{ijkl}. \quad [8]$$

The main effect of condition tested whether intrapersonal recurrence differed across listening conditions after accounting for variation across the signed lag profile. Lag bin was entered as a categorical predictor so that no linear or symmetric form of the DCRP was assumed. Tukey-adjusted estimated marginal mean comparisons were used to compare recurrence between *Noise-Self*, *Noise-Other*, and *Noise-Free*, averaged across lag bins.

### References


1. H He, HV Skat-Rørdam, I Örnólfsson, I Konvalinka, An automated pipeline for standardised speech-unit annotation in spontaneous dialogue (2026).
2. S Wallot, Multidimensional cross-recurrence quantification analysis (mdcrqa)–a method for quantifying correlation between multivariate time-series. *Multivar. behavioral research* **54**, 173–191 (2019).
3. DC Richardson, R Dale, Looking to understand: The coupling between speakers' and listeners' eye movements and its relationship to discourse comprehension. *Cogn. science* **29**, 1045–1060 (2005).
